# Polynuclear ruthenium amines inhibit K_2P_ channels via a ‘finger in the dam’ mechanism

**DOI:** 10.1101/863837

**Authors:** Lianne Pope, Marco Lolicato, Daniel L. Minor

**Author notes:** Department of Molecular Medicine, University of Pavia, Pavia ITALY (Marco Lolicato).

## Abstract

The trinuclear ruthenium amine Ruthenium Red (RuR) inhibits diverse ion channels including K_2P_ potassium channels, TRPs, the mitochondrial calcium uniporter, CALHMs, ryanodine receptors, and Piezos. Despite this extraordinary array, there is very limited information for how RuR engages its targets. Here, using X-ray crystallographic and electrophysiological studies of an RuR-sensitive K_2P_, K_2P_2.1 (TREK-1) I110D, we show that RuR acts by binding an acidic residue pair comprising the ‘Keystone inhibitor site’ under the K_2P_ CAP domain archway above the channel pore. We further establish that Ru360, a dinuclear ruthenium amine not known to affect K_2P_s, inhibits RuR-sensitive K_2P_s using the same mechanism. Structural knowledge enabled a generalizable RuR ‘super-responder’ design strategy for creating K_2P_s having nanomolar sensitivity. Together, the data define a ‘finger in the dam’ inhibition mechanism acting at a novel K_2P_ inhibitor binding site. These findings highlight the polysite nature of K_2P_ pharmacology and provide a new framework for K_2P_ inhibitor development.

## Introduction

Ruthenium red (RuR) (Fletcher et al., 1961) (Figure 1A) is a trinuclear oxo-bridged ruthenium amine polycation with many biological applications (Clarke, 2002), including a ∼50 year legacy of use as an inhibitor of diverse ion channels, such as select members of the K_2P_ (KCNK) family (Braun et al., 2015; Czirjak and Enyedi, 2003; Gonzalez et al., 2013; Musset et al., 2006), numerous TRP channels (Arif Pavel et al., 2016; Caterina et al., 1999; Caterina et al., 1997; Guler et al., 2002; Story et al., 2003; Strotmann et al., 2000; Voets et al., 2004; Voets et al., 2002), the mitochondrial calcium uniporter (MCU) (Chaudhuri et al., 2013; Kirichok et al., 2004; Moore, 1971; Rahamimoff and Alnaes, 1973), CALHM calcium channels (Choi et al., 2019; Dreses-Werringloer et al., 2013; Ma et al., 2012), ryanodine receptors (Ma, 1993; Smith et al., 1988), and Piezo channels (Coste et al., 2012; Zhao et al., 2016). Despite this remarkably wide range of ion channel targets and the recent boom in ion channel structural biology, structural understanding of how RuR acts on any ion channel is limited to a recent cryo-EM study of the CALHM2 channel that provides few molecular details regarding the coordination chemistry that underlies RuR binding (Choi et al., 2019). In the case of K_2P_s, functional studies have established that a negatively charged residue at the base of the K_2P_ extracellular domain that forms an archway over the channel pore, the CAP domain, comprises a key RuR sensitivity determinant in the natively RuR sensitive channels K_2P_9.1 (TASK-3) (Czirjak and Enyedi, 2003; Gonzalez et al., 2013; Musset et al., 2006) and K_2P_10.1 (TREK-2) (Braun et al., 2015). Further, installation of a negatively charged amino acid at the equivalent CAP domain site in a non-RuR sensitive channel is sufficient to confer RuR sensitivity (Braun et al., 2015). The archway above the selectivity filter extracellular mouth made by the K_2P_ CAP domain creates a pair of water-filled portals, the extracellular ion pathway (EIP), through which ions exit the channel under physiological conditions (Brohawn et al., 2012; Dong et al., 2015; Lolicato et al., 2017; Miller and Long, 2012). Although the EIP has been proposed as the site of RuR action (Braun et al., 2015; Gonzalez et al., 2013), the mechanism by which RuR inhibits K_2P_s remains unresolved and to date there is no direct structural evidence indicating that the EIP can be targeted by RuR or any other class of small molecule or protein-based inhibitors.

**Figure 1.**
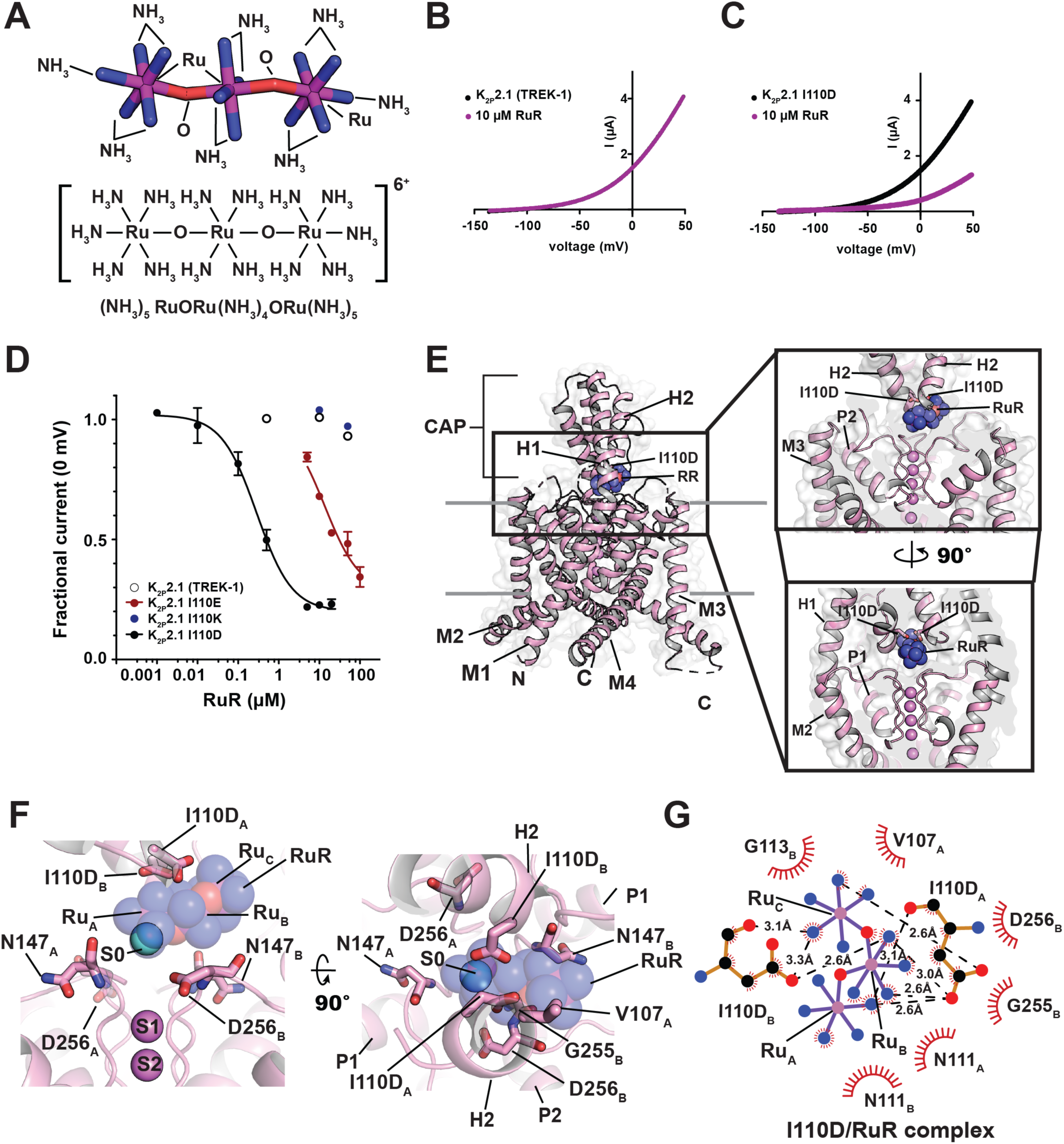
Functional and structural analysis of the K_2P_2.1 I110D:RuR complex. (**A**) Ruthenium Red (RuR) structure (**B**) and (**C**) Exemplar TEVC recordings of (**B**) K_2P_2.1 (TREK-1) and (**C**) K_2P_2.1 I110D responses to 10 µM RuR (magenta). (**D**) Dose-response curves for K_2P_2.1 (TREK-1) (open white circles), K_2P_2.1 I110D (black), K_2P_2.1 I110E (red), and K_2P_2.1 I110K to RuR. (**E**) Structure of the K_2P_2.1 I110D:RuR complex. Inset shows the location of the RuR binding site. I110D is shown as sticks. (**F**) Close up view of K_2P_2.1 I110D:RuR interactions. S0 ion (cyan) from the K_2P_2.1 I110D structure is indicated, RuR is shown in space filling in panels (**E**) and (**F**). (**G**) LigPLOT (Wallace et al., 1995) diagram of K_2P_2.1 I110D:RuR interactions showing ionic interactions (dashed lines) and van der Waals contacts (red) ≤5Å.

K_2P_s produce an outward ‘leak’ potassium current that plays a critical role in stabilizing the resting membrane potential of diverse cell types in the nervous, cardiovascular, and immune systems (Enyedi and Czirjak, 2010; Feliciangeli et al., 2014; Renigunta et al., 2015). There are fifteen K_2P_ subtypes comprising six subfamilies in which the channel monomers assemble into dimers wherein each subunit contributes two conserved pore forming domains to make the channel pore (Brohawn et al., 2012; Dong et al., 2015; Feliciangeli et al., 2014; Lolicato et al., 2017; Miller and Long, 2012; Rödström et al., 2019). A range of physical and chemical signals control K_2P_ function (Enyedi and Czirjak, 2010; Feliciangeli et al., 2014; Renigunta et al., 2015) and various K_2P_ subtypes have emerging roles in a multitude of physiological responses and pathological conditions such as action potential propagation in myelinated axons (Brohawn et al., 2019; Kanda et al., 2019), anesthetic responses (Heurteaux et al., 2004; Lazarenko et al., 2010), microglial surveillance (Madry et al., 2018), sleep duration (Yoshida et al., 2018), pain (Alloui et al., 2006; Devilliers et al., 2013; Vivier et al., 2017), arrythmia (Decher et al., 2017), ischemia (Heurteaux et al., 2004; Laigle et al., 2012; Wu et al., 2013), cardiac fibrosis (Abraham et al., 2018), depression (Heurteaux et al., 2006), migraine (Royal et al., 2019), intraocular pressure regulation (Yarishkin et al., 2018), and pulmonary hypertension (Lambert et al., 2018). Although there have been recent advances in identifying new K_2P_ modulators (Bagriantsev et al., 2013; Lolicato et al., 2017; Pope et al., 2018; Su et al., 2016; Tian et al., 2019; Vivier et al., 2017; Wright et al., 2019) and in defining key structural aspects of K_2P_ channel pharmacology (Dong et al., 2015; Lolicato et al., 2017; Schewe et al., 2019), as is the case with many ion channel classes, pharmacological agents targeting K_2P_s remain poorly developed and limit the ability to probe K_2P_ mechanism and biological functions (Sterbuleac, 2019).

Here, we present X-ray crystal structures of a K_2P_2.1 (TREK-1) mutant bearing a single change at the site that controls K_2P_ channel RuR sensitivity, K_2P_2.1 I110D, alone and complexed with two different polynuclear ruthenium amines, RuR and the dinuclear ruthenium amine, Ru360 (Ying et al., 1991), an inhibitor of the mitochondrial calcium uniporter (Baughman et al., 2011; Kirichok et al., 2004; Oxenoid et al., 2016) not previously known to affect potassium channels. The structures show that the negatively charged residues at position 110 comprise ‘the Keystone inhibitor site’ on the ceiling of the CAP archway to which positively charged RuR and Ru360 bind through ionic interactions. This interaction holds the polybasic compounds directly over the mouth of the channel pore, blocks one EIP arm, and prevents channel function. Functional studies corroborated by a crystal structure of K_2P_2.s I110D bound simultaneously to RuR and a small molecule activator of the channel selectivity filter ‘C-type gate’, ML335 (Lolicato et al., 2017), establish that polynuclear ruthenium amine inhibition of K_2P_s is unaffected by C-type gate activation. Using molecular recognition principles derived from the structures of the K_2P_:RuR and K_2P_:Ru360 complexes, we demonstrate a general design strategy for endowing any K_2P_ channel with nanomolar RuR sensitivity. Our work establishes that polynuclear ruthenium compounds act through a ‘finger in the dam’ mechanism to inhibit K_2P_ function by binding under the CAP domain archway at the Keystone inhibitor site and blocking the pore. The structural definition of this new modulatory site demonstrates the importance of electronegativity and specific sidechain geometry for polynunclear amine molecular recognition, defines a new small molecule binding site that augments the rich, polysite pharmacology of K_2P_ modulation, and opens a path for targeting the Keystone inhibitor site and EIP for the development of new K_2P_ modulators.

## Results

### A single site in the K_2P_ CAP domain confers RuR sensitivity

K_2P_2.1 (TREK-1) is the founding member of the thermo- and mechanosensitive subgroup of K_2P_s (Douguet and Honore, 2019; Feliciangeli et al., 2014). Although this channel is resistant to RuR inhibition (Figure 1B and D, Table 1) (Braun et al., 2015), two electrode voltage clamp (TEVC) recordings in *Xenopus* oocytes of outward current inhibition by RuR under physiological ionic conditions showed that installation of a point mutation I110D at the base of the K_2P_2.1 (TREK-1) CAP domain conferred sub-micromolar RuR sensitivity to K_2P_2.1 (TREK-1) (IC_50_ = 0.287 ± 0.054 µM)(Figs. 1C-D, Table 1). This inhibition followed a 1:1 RuR:channel stoichiometry, in agreement with K_2P_ studies using other recording protocols (Braun et al., 2015; Czirjak and Enyedi, 2003), and validates previous studies showing that this point mutant renders K_2P_2.1 (TREK-1) sensitive to RuR (Braun et al., 2015). Importantly, the response of K_2P_2.1 I110D to RuR matched that of the closely-related, natively-RuR sensitive K_2P_10.1 (TREK-2) (Braun et al., 2015) in which there are native aspartate residues at the K_2P_2.1 Ile110 analogous site (Figs. 1D and S1A, Table 1) (IC_50_ = 0.287 ± 0.054 and 0.23 ± 0.06 µM for K_2P_2.1 I110D and K_2P_10.1 (TREK-2) (Braun et al., 2015), respectively), suggesting that the I110D change to K_2P_2.1 (TREK-1) captures the essence of the requirements for RuR inhibition.

**Table 1.**
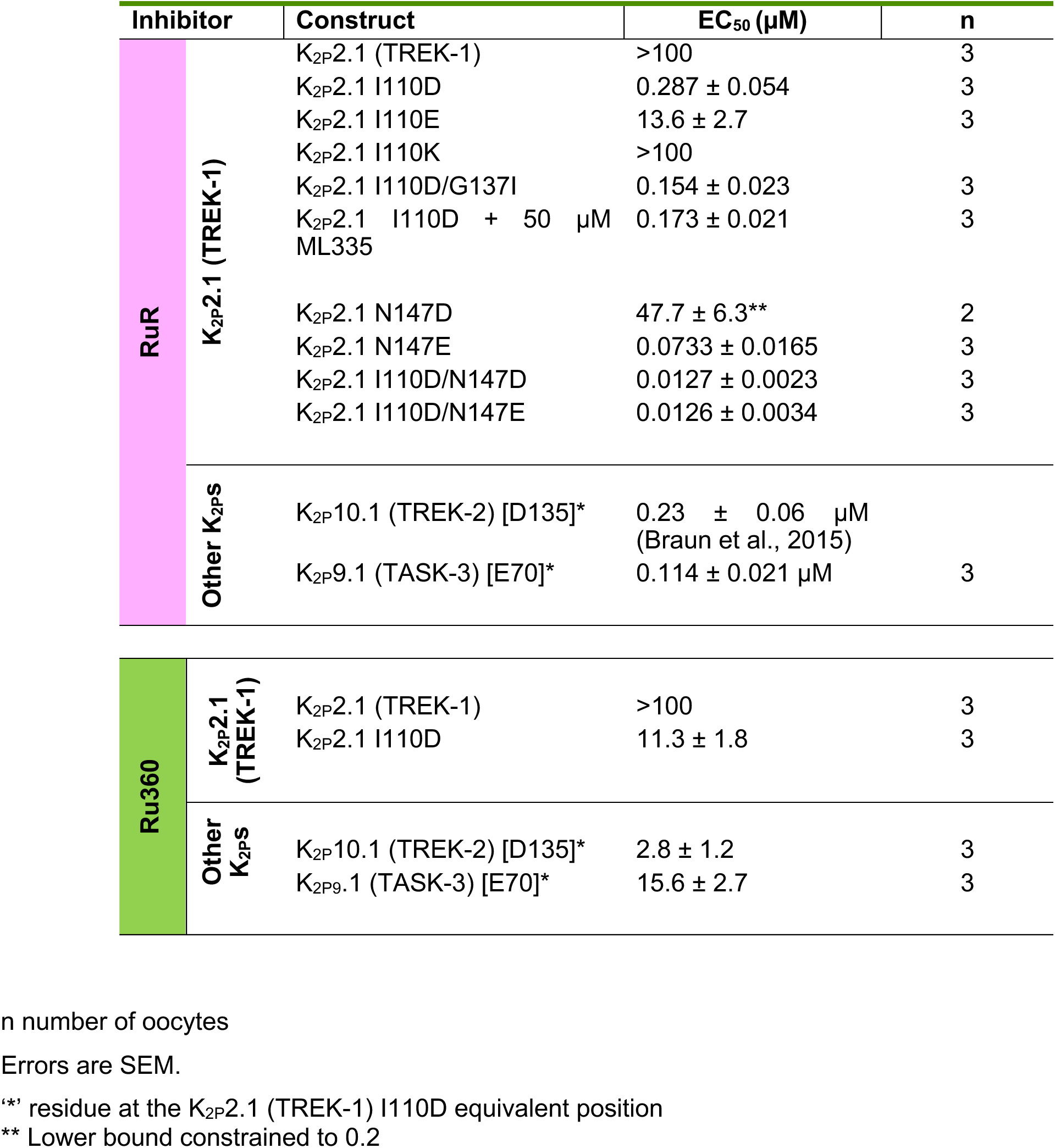
IC_50_ values for RuR and Ru360.

Because another natively-RuR sensitive K_2P_, K_2P_9.1 (TASK-3), has a glutamate at the K_2P_2.1 I110D equivalent site (Figure S1A) that is essential for its RuR response (Czirjak and Enyedi, 2003; Gonzalez et al., 2013; Musset et al., 2006), we asked whether I110E would also render K_2P_2.1 (TREK-1) sensitive to RuR. Indeed, TEVC measurements showed that RuR inhibited K_2P_2.1 I110E (IC_50_ =13.6 ± 2.7 µM, Table 1) (Figs. 1D and S1B, Table 1). RuR inhibition was ∼50 fold weaker than that observed for K_2P_2.1 I110D, indicating that the I110D and I110E changes are not equivalent even though both bear similar negative charges. Further, introduction of a positively charged residue, K_2P_2.1 I110K, yielded channels as insensitive to RuR as K_2P_2.1 (TREK-1) (Figs.1D and S1C). RuR inhibition was essentially independent of voltage for K_2P_2.1 I110D and K_2P_2.1 I110E (Figs. S1D-G), consistent with prior reports of RuR inhibition of other K_2P_s (Czirjak and Enyedi, 2002). Together, these results provide key support for the idea that a negative charge at the K_2P_ CAP domain base is a crucial determinant of RuR inhibition of K_2P_s (Braun et al., 2015; Czirjak and Enyedi, 2003; Gonzalez et al., 2013; Musset et al., 2006). Importantly, the observation of the ∼50 fold difference between the RuR sensitivity of two essentially equivalently negatively charged residues in the same structural context, K_2P_2.1 I110D and K_2P_2.1 I110E (Figure 1D, Table 1) indicates that electrostatics is not the sole factor contributing to the RuR:channel interaction and points to a role for the detailed geometry of the interaction of the negatively charged residues with RuR.

### Structural definition of the K_2P_ RuR binding site

To understand how RuR inhibits K_2P_ channels, we determined the X-ray crystal structures of K_2P_2.1 I110D alone and bound to RuR at resolutions of 3.40Å and 3.49Å, respectively (Table S1) on the background of the previously crystallized construct K_2P_2.1 (TREK-1)_cryst_ (Lolicato et al., 2017). Apart from the I110D change, the overall structure of K_2P_2.1 I110D was essentially identical to K_2P_2.1 (TREK-1)_cryst_ (RMSD_Cα_ = 0.575Å). Importantly, K_2P_2.1 I110D was structurally similar to the natively-RuR sensitive K_2P_10.1 (TREK-2) (Dong et al., 2015) (RMSD_Cα_= 0.938Å), especially in the neighborhood of K_2P_10.1 (TREK-2) Asp140 (Figure S2A, Table S2), the residue that is fundamental to K_2P_10.1 (TREK-2) RuR sensitivity (Braun et al., 2015). Hence, when taken together with the functional similarity to K_2P_10.1 (TREK-2), the K_2P_2.1 I110D:RuR complex should capture the essential elements that contribute to the RuR response of natively RuR-sensitive K_2P_s.

The K_2P_2.1 I110D:RuR complex structure shows that RuR binds under the CAP domain archway directly above the selectivity filter at a site we term the ‘Keystone inhibitor site’ due to the location of the I110D residues at the peak of the CAP archway ceiling (Figs. 1E and S2B-C). RuR binds with a 1:1 stoichiometry to the channel that matches expectations from functional studies (Table 1) (Braun et al., 2015; Czirjak and Enyedi, 2002; 2003). To facilitate description of this and other RuR complexes, we designate the three RuR ruthenium amine centers as Ru_A_, Ru_B_, and Ru_C_. RuR binds at ∼45° angle relative to the CAP and selectivity filter in a pose that places one of the terminal ruthenium-amine moieties, Ru_A_, directly above the column of selectivity filter ions in a position that overlaps with the S0 ion site from the K_2P_2.1 I110D structure (Figs. 1F and S2B). The Ru_B_ and Ru_C_ moieties block one EIP arm (Figs. 1E-F). Notably, the observed pose is very different from the previously proposed horizontal RuR binding pose in which the Ru_B_ moiety sits above the column of selectivity filter ions (Gonzalez et al., 2013). The I110D carboxylates coordinate RuR directly through electrostatic interactions with all three ruthenium-amine moieties using a multipronged set of interactions (Figure 1F and G). Such direct coordination suggests why K_2P_2.1 I110D and K_2P_2.1 I110E have different magnitude RuR responses, as the extra methylene groups in K_2P_2.1 I110E would not allow the same type of direct coordination observed for the smaller aspartate pair (Figure 1G). Besides direct electrostatic interactions, there are van der Waals contacts between Ru_C_ and the side chain of CAP residue Val107 from chain A, Ru_A_ and Ru_B_ with Asn111 from each chain of the dimer, Ru_A_ with Asp256 from chain B, and Ru_B_ with Gly255 from chain B. Apart from a slight reorientation of the I110D sidechains and CAP to accommodate RuR (Figure S2D, Movie S1), there are only minor conformational changes with respect to the unbound K_2P_2.1 I110D (RMSD_Cα_= 0.688Å)(Table S2). Hence, RuR binds to an essentially pre-organized electronegative binding site at the CAP base.

### K_2P_ RuR binding is independent of C-type gate activation

Activation of the selectivity filter, ‘C-type’ gate is central to K_2P_ function (Bagriantsev et al., 2012; Bagriantsev et al., 2011; Lolicato et al., 2017; Piechotta et al., 2011; Schewe et al., 2016). Because RuR binds directly above the selectivity filter and overlaps with the S0 ion, we asked whether C-type gate activation by mutation, G137I (Bagriantsev et al., 2012; Lolicato et al., 2014) or by a small molecule activator, ML335 (Lolicato et al., 2017), would impact RuR inhibition of K_2P_2.1 I110D (Figure 2A-B). TEVC experiments showed that RuR inhibits K_2P_2.1 I110D/G137I with an IC_50_ (IC_50_ = 0.154 ± 0.023 µM) that is very similar to that for K_2P_2. I110D, indicating that C-type gate activation does not influence RuR block of the channel (Figure 2C, Table 1).

**Figure 2.**
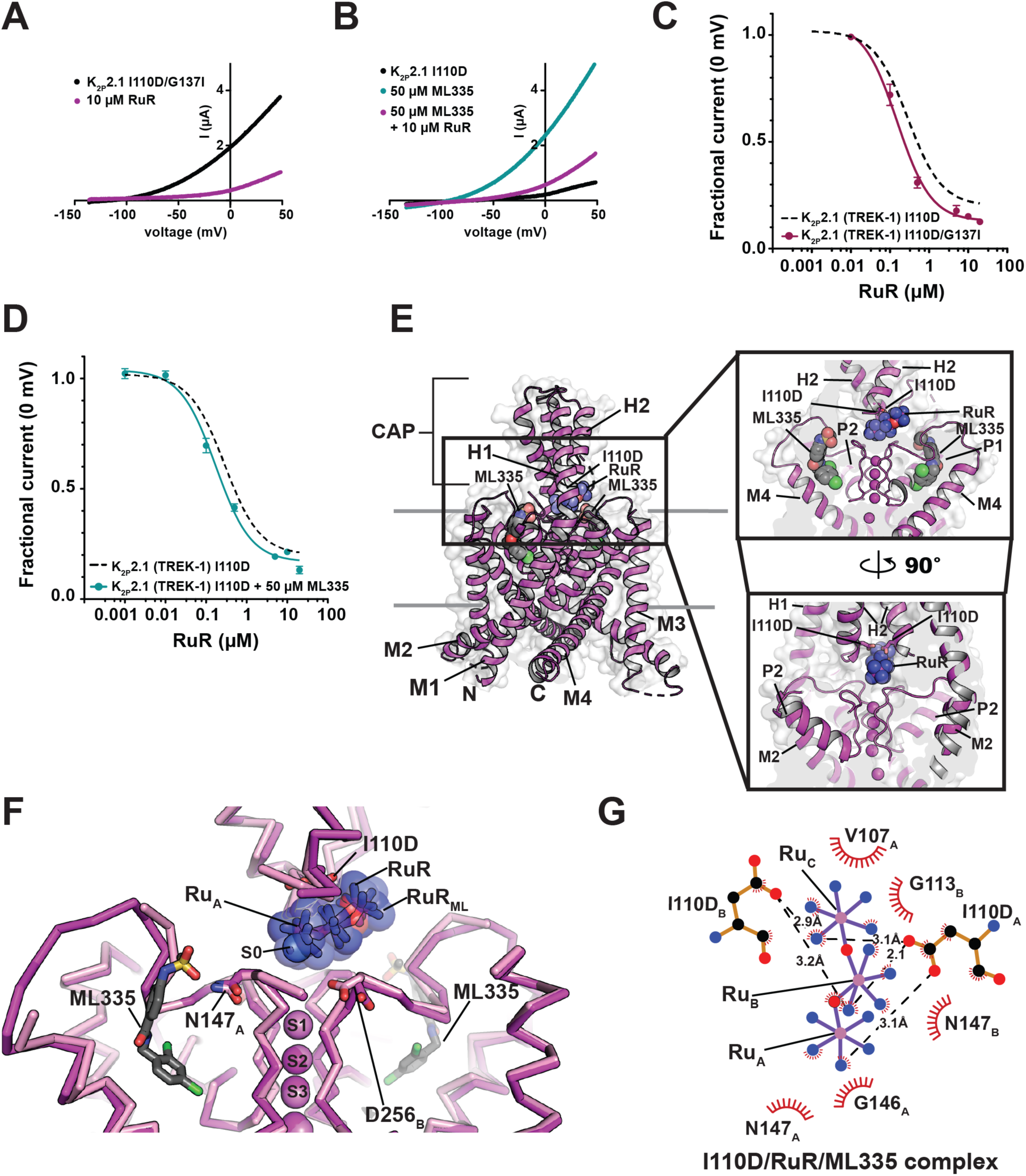
Functional and structural analysis of C-type gate activated K_2P_2.1 I110D:RuR complexes. (**A**) and (**B**) Exemplar TEVC recordings of: (**A**) K_2P_2.1 I110D/G137I (black) and in the presence of 10 µM RuR (magenta), and (**B**) K_2P_2.1 I110D alone (black) in the presence of 50µM ML335 (cyan), and in the presence of 50 µM ML335 +10 µM RuR (magenta). (**C**) and (**D**) RuR dose-response curves for (**C**) K_2P_2.1 I110D/G137I (magenta), and (**D**) K_2P_2.1 I110D in the presence of 50 µM ML335 (cyan). Dashed lines show RuR response of K_2P_2.1 I110D from Figure 1D. (**E**) Structure of the K_2P_2.1 I110D:ML335 RuR complex. Inset shows the location of the RuR binding site. I110D is shown as sticks. ML335 (grey) is shown as space filling. (**F**) Superposition of RuR binding site from the K_2P_2.1 I110D:RuR (pink) and K_2P_2.1 I110D:ML335:RuR (magenta) RuR and RuR_ML_ indicate RuR from the K_2P_2.1 (TREK-1) I110D:RuR and K_2P_2.1 I110D:ML335:RuR structures, respectively. ML335 is shown as sticks. (**G**) LigPLOT (Wallace et al., 1995) diagram of K_2P_2.1 I110D:RuR interactions from the K_2P_2.1 I110D:ML335:RuR complex showing ionic interactions (dashed lines) and van der Waals contacts (red) ≤5Å.

Before assessing whether RuR inhibition was influenced by ML335 activation, we first measured the activation of K_2P_2.1 I110D by ML335. The I110D change is >15Å from the K_2P_ modulator pocket that forms the ML335 binding site and is not expected to impact ML335 activation. In line with these expectations, there was no difference in the response of K_2P_2.1 I110D to ML335 activation relative to K_2P_2.1 (TREK-1) (EC_50 ML335)_ = 11.8 ± 2.3 and 14.3 ± 2.7 µM for K_2P_2. I110D and K_2P_2.1 (TREK-1) (Lolicato et al., 2017), respectively) (Figs. S3A-B). Importantly, similar to the observations with the G137I C-type gate activation mutant, pharmacological C-type gate activation by saturating amounts of ML335 had minimal impact on the RuR response relative to K_2P_2.1 I110D (IC_50_ = 0.173 ± 0.021 µM, Table 1). Together, these data demonstrate that RuR inhibition is essentially independent of C-type gate activation (Figs. 2A-D, Table 1) and are consistent with the observed position of RuR above the selectivity filter.

To see whether there might be structural differences in the interaction of RuR with an activated C-type gate, we determined the structure of the K_2P_2.1 I110D:RuR:ML335 complex at 3.00Å resolution (Figs. 2E and S3C-D, Table S1). This structure is very similar to the K_2P_2.1 I110D:RuR complex (Figs. S3C-E) (RMSD_Cα_ = 0.507Å) and to the previously determined K_2P_2.1:ML335 complex (Lolicato et al., 2017) (RMSD_Cα_ = 0.480Å) (Table S2). The structure shows that RuR binds to the Keystone inhibitor site using a pose that is very similar to that in the K_2P_2.1 I110D:RuR complex. Both I110D sidechains coordinate multiple Ru centers though direct interactions to RuR (Figure 2F-G) like those in the K_2P_2.1 I110D:RuR complex and there are van der Waals contacts with residues in the CAP and selectivity filter outer mouth. These fundamental similarities in binding to the Keystone inhibitor site are consistent with the similar IC_50_s measured with or without ML335 C-type gate activation (Figure 2C-D,Table 1).

### Polynuclear ruthenium compounds inhibit K_2P_s at a common site

The dinuclear oxo-bridged ruthenium compound, Ru360 (Figure 3A) (Ying et al., 1991), has many characteristics in common with RuR (cf. Figure 1A). Ru360 is best known as an inhibitor of the mitochondrial calcium uniporter (MCU) (Baughman et al., 2011; Kirichok et al., 2004; Oxenoid et al., 2016), a property it shares with RuR (Gunter and Pfeiffer, 1990). Yet, despite its structural similarity to RuR and the fact that both Ru360 and RuR inhibit MCU, Ru360 has not been reported to block K_2P_ channels. To ask whether Ru360 might inhibit RuR-sensitive K_2P_ channels, we used TEVC to measure the Ru360 responses of K_2P_2.1 (TREK-1), K_2P_2.1 I110D, and the natively RuR sensitive K_2P_s, K_2P_10.1 (TREK-2) (Braun et al., 2015) and K_2P_9.1 (TASK-3) (Czirjak and Enyedi, 2003; Gonzalez et al., 2013; Musset et al., 2006). Application of 100 µM Ru360 to K_2P_2.1 (TREK-1) had no effect, consistent with the insensitivity of this channel to RuR (Figs. 3B and F). However, in stark contrast, Ru360 inhibited all of the RuR sensitive K_2P_s with micromolar potency (IC_50_ = 11.3 ± 1.8, 2.8 ± 1.2, and 15.6 ± 2.7 µM for K_2P_2.1 I110D, K_2P_10.1 (TREK-2), and K_2P_9.1 (TASK-3), respectively) (Figs. 3C-F, Table 1). Similar to RuR inhibition, Ru360 block of K_2P_s was independent of voltage (Figs. S4A-D). For both K_2P_2.1 I110D and K_2P_10.1 (TREK-2) the Ru360 IC_50_s are >10-fold weaker than those for RuR. Because there is a reported 30-fold discrepancy in the IC_50_ of RuR for K_2P_9.1 (TASK-3) in the literature (0.35 µM (Czirjak and Enyedi, 2003) vs.10 µM (Gonzalez et al., 2013)) that precluded a direct comparison with our Ru360 data, we measured inhibition of K_2P_9.1 (TASK-3) by RuR to resolve whether RuR and Ru360 had similar or different IC_50_s for K_2P_9.1 (TASK-3). Our TEVC experiments measured a sub-micromolar IC_50_ for RuR inhibition of K_2P_9.1 (TASK-3) (IC_50_ = 0.114 ± 0.021 µM) (Figs. S4E-F, Table 1) that agrees with other *Xenopus* oocyte TEVC studies (Czirjak and Enyedi, 2003). These data establish that, as with the other polynuclear ruthenium-sensitive K_2P_s we studied, Ru360 is a weaker inhibitor of K_2P_9.1 (TASK-3) than RuR. The uniformly weaker potency of Ru360 versus RuR against K_2P_s correlates with the fact that Ru360 carries half the positive charge of RuR (+3 versus +6) and underscores the important role that electrostatics plays in the binding of these polycations.

**Figure 3.**
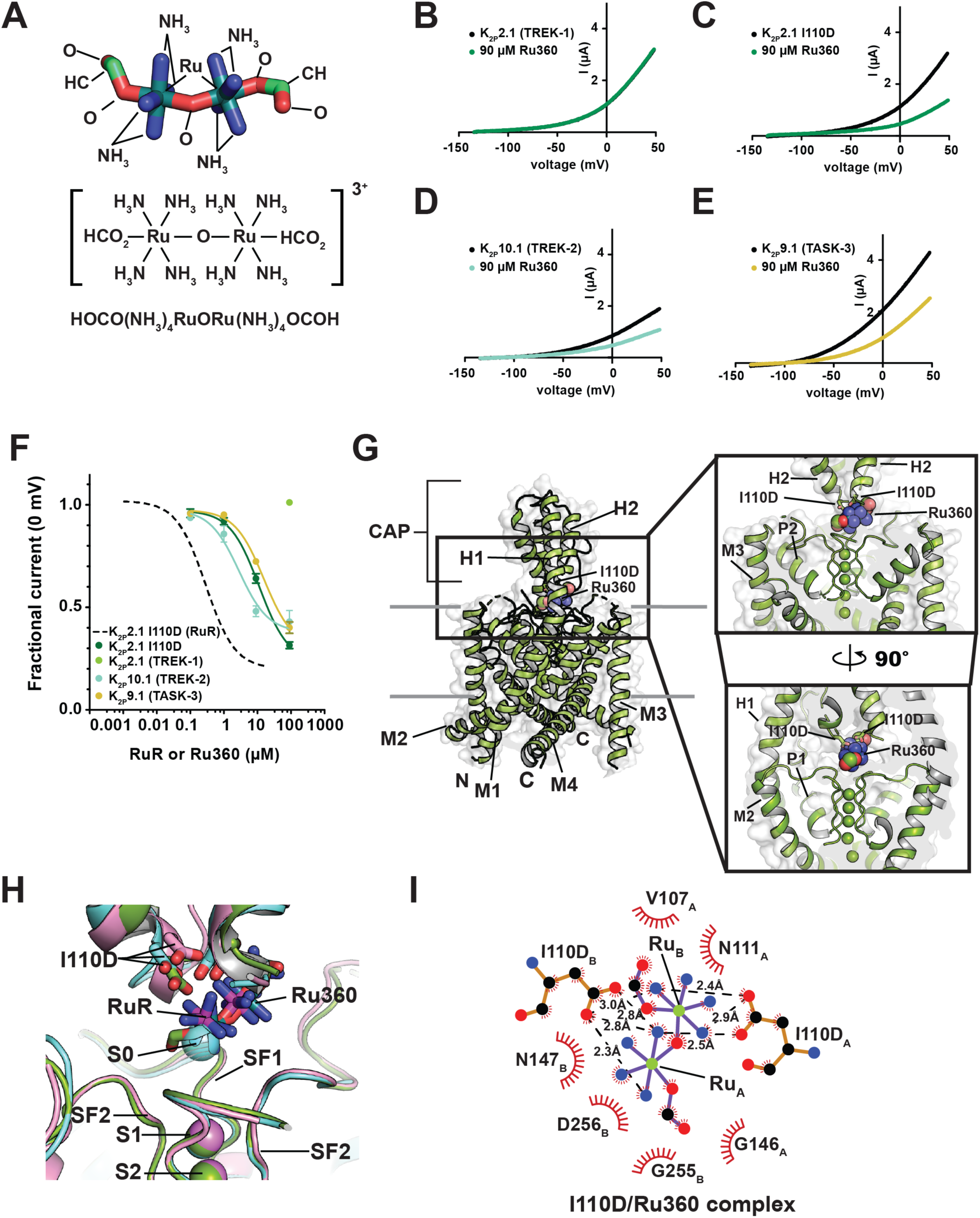
Functional and structural analysis of K_2P_:Ru360: interactions. (**A**) Ru360 (RuR) structure. (**B-D**) Exemplar TEVC recordings of (**B**) K_2P_2.1 (TREK-1), (**C**) K_2P_2.1 I110D, (**D**) K_2P_10.1 (TREK-2), and (**E**) K_2P_9.1 (TASK-3) alone (black) and in the presence of 90 µM Ru360 (green, teal, and olive, respectively). (**F**) Ru360 dose-response curves for K_2P_2.1 I110D (green), K_2P_2.1 (TREK-1) (light green), K_2P_10.1 (TREK-2) (teal), and K_2P_9.1 (TASK-3) (olive). Dashed line shows RuR dose-response for K_2P_2.1 I110D from Figure 1D. (**G**) Structure of the K_2P_2.1 I110D:Ru360 complex. Inset shows the location of the Ru360 binding site. I110D is shown as sticks. (**H**) Close up view of K_2P_2.1 I110D (cyan), K_2P_2.1 I110D:RuR (pink), and K_2P_2.1 I110D:Ru360 Keystone inhibitor sites. RuR and Ru360 are shown as sticks. S0 ion from the K_2P_2.1 I110D structure is shown as a sphere. (**I**) LigPLOT (Wallace et al., 1995) diagram of K_2P_2.1 I110D:Ru360 interactions showing ionic interactions (dashed lines) and van der Waals contacts (red) ≤5Å.

To understand the details of how Ru360 inhibits K_2P_s, we determined of a 3.51Å resolution X-ray crystal structure of the K_2P_2.1 I110D:Ru360 complex (Figs. 3G-H, Table S1). As with the RuR complexes, the K_2P_2.1 I110D:Ru360 complex has a channel structure that is overall very similar to the structure of the K_2P_2.1 I110D in the absence of the inhibitor (RMSD_Cα_ = 0.665Å) and to the K_2P_2.1 I110D:RuR complex (RMSD_Cα_ = 0.561Å) (Figure S4G, Table S2). Ru360 binds to the Keystone inhibitor site in a pose that matches RuR (Figs. 3H and S4G-I). Notably, even though Ru360 has one fewer ruthenium atoms than RuR, one of the Ru360 ruthenium-amine moieties, denoted Ru_A_, occupies essentially the same site as the RuR Ru_A_ moiety and overlaps with the S0 ion site (Figure 3H), while the other ruthenium amine, Ru_B_, overlaps the position of the RuR Ru_B_ moiety and blocks one EIP arm. There are essentially no conformational changes between K_2P_2.1 I110D and the K_2P_2.1 I110D:Ru360 complex except for a change near the base of the CAP helices similar to that seen in the RuR complexes (Figure S4I). The Keystone inhibitor site acidic sidechains directly coordinate the Ru360 Ru_A_ and Ru_B_ centers (Figure 3I). Ru360 also makes van der Waals contacts to the upper part of the selectivity filter and part of the CAP from chain A similar to those made by RuR (cf. Figs. 1G, 2G, and 3I). Together, these data demonstrate that Ru360 inhibits RuR-sensitive K_2P_s and reveals the common mode by which polynuclear ruthenium amines affect K_2P_ channels by binding to the Keystone inhibitor site, blocking ion exit from the selectivity filter, and obstructing one EIP arm.

### Protein engineering creates RuR super-responders

Because the electronegative nature of the Keystone inhibitor site is a key determinant of K_2P_ channel sensitivity to polynuclear ruthenium compounds, we wanted to test whether increasing the electronegative character of surrounding portions of the EIP would also affect RuR block. To identify candidate sites, we looked for elements on the floor of the RuR and Ru360 binding site that made close contacts with the inhibitors. This analysis identified the backbone atoms of the selectivity filter outer mouth residues Asn147 and Asp256 as the nearest neighbors (Figs. 1G, 2G, 3G, and 4A). As we cannot easily change the backbone atoms, we considered changing the properties of the sidechains from these positions. Sequence comparison of representatives from each K_2P_ subtype (Figure S5A) shows that the two sites have very different conservation patterns. The Asn147 site shows a range of amino acid types. This variability contrasts with the strict conservation at the Asp256 site. Because all K_2P_s have aspartate at the 256 position regardless of whether or not they RuR-sensitive, we reasoned that the negatively charged sidechain at this site has no influence on RuR binding. By contrast, the amino acid diversity at the Asn147 site indicated that this site might have different properties than the Asp256 site. Hence, we tested whether replacing Asn147 with a negatively charged residue would impact K_2P_2.1 (TREK-1) RuR sensitivity. TEVC experiments showed that RuR inhibited both K_2P_2.1 N147D and K_2P_2.1 N147E (Figs. 4C-F), demonstrating that the presence of an acidic residue at the Keystone inhibitor site is not the only means by which a K_2P_ channel can acquire RuR sensitivity. Notably, there was a marked difference in the IC_50_s between the two mutants with N147E having a much greater susceptibility to RuR inhibition than N147D (IC_50_ = 0.0733 ± 0.0165 and 47.7 ± 6.3 µM for K_2P_2.1 N147E and K_2P_2.1 N147D, respectively) (Figs. 4G-I, Table 1). Similar to RuR inhibition of other K_2P_s, RuR block of K_2P_2.1 N147E was essentially voltage independent, whereas K_2P_2.1 N147D showed a mild voltage-dependence (Figs. S5B-E).

**Figure 4.**
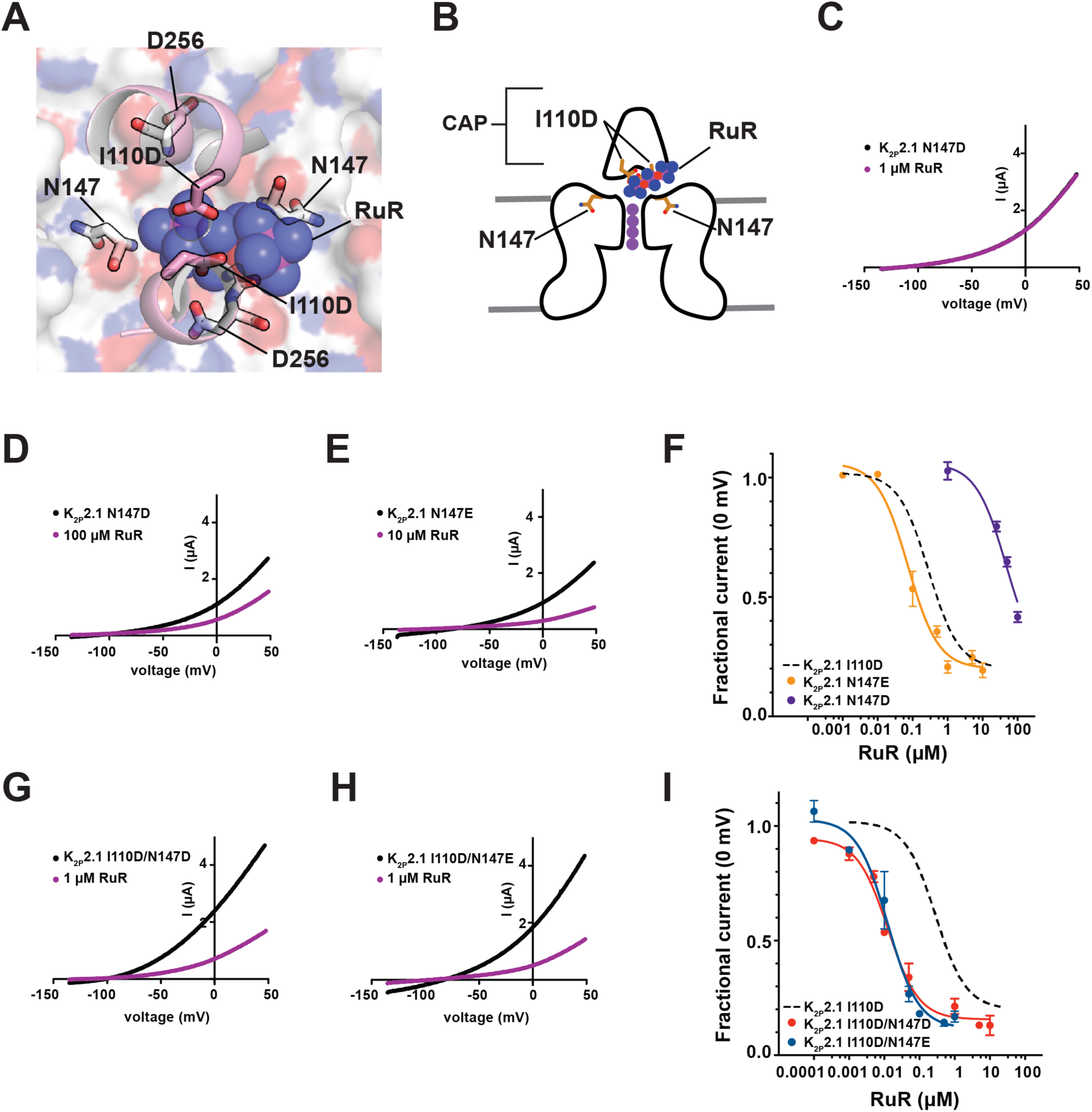
Engineering K_2P_ RuR super-responders. (**A**) View through the K_2P_2.1 (TREK-1) CAP to the floor of the Keystone inhibitor site. CAP H2 helix (pink) is shown as a cartoon. RuR is shown as semi-transparent spheres. Surface (white) shows the top of the selectivity filter. I110D, N147, and D256 are shown as sticks. (**B**) Cartoon depiction of the elements framing the Keystone inhibitor site. Locations of CAP, I110D, and N147 are indicated. Selectivity filter potassium ions are shown as purple circles. (**C-E**) TEVC recordings of (**C**) K_2P_2.1 N147D alone (black) and in the presence of 1 µM RuR (magenta), (**D**) K_2P_2.1 N147D alone (black) and in the presence of 100 µM RuR (magenta) and (**E**) K_2P_2.1 N147E alone (black) and in the presence of 10 µM RuR (magenta). (**F**) RuR response of K_2P_2.1 N147D (purple) and K_2P_2.1 N147E (orange). Dashed line shows K_2P_2.1 I110D response to RuR, from Figure 1D. (**G**) and (**H**) TEVC recordings of (**G**) K_2P_2.1 I110D/N147D alone (black) and in the presence of 1 µM RuR (magenta), and (**H**) K_2P_2.1 I110D/N147E alone (black) and in the presence of 1 µM RuR (magenta). (**I**) RuR dose-response of K_2P_2.1 I110D/N147D (red) and K_2P_2.1 I110D/N147E (blue). Dashed line shows K_2P_2.1 I110D response to RuR, from Figure 1D.

Because both N147D and N147E changes were able to confer RuR sensitivity to K_2P_2.1 (TREK-1) (Figure 4F, Table 1), we asked whether having negatively charged residues on both the ceiling (residue 110) and floor (residue 147) of the RuR binding site would result in enhanced RuR inhibition. TEVC measurements of the RuR responses of K_2P_2.1 I110D/N147D and K_2P_2.1 I110D/N147E revealed that both double mutants had similar IC_50_s (IC_50_= 0.0127 ± 0.0023 and 0.0126 ± 0.0034 µM for K_2P_2.1 I110D/N147D and K_2P_2.1 I110D/N147E, respectively) (Figs. S5F-G, Table 1). Similar to the I110D and N147E mutants, the response of both double mutant channels to RuR was essentially independent of voltage (Figs. S5H-J).

The IC_50_s values of the double mutants were an order of magnitude better than that of K_2P_2.1 I110D alone and three orders of magnitude better than K_2P_2.1 N147D. Hence, we turned to double mutant cycle analysis (Carter et al., 1984; Hidalgo and MacKinnon, 1995) to assess the extent of synergy between the two sites with respect to RuR inhibition. This analysis uncovered a strong positive cooperativity for the I110D/N147D pair (ΔΔG = −4.1 kcal mol^-1^) (Figure S6A). By contrast, the enhanced RuR response of the I110D/N147E combination resulted from essentially additive contributions of the two negatively charged residues (ΔΔG = −0.3 kcal mol^-1^) (Figure S6B). The fact that the two double mutant pairs do not behave equivalently even though both comprise two sets of acidic sidechains, together with the observation that there is a substantial difference in the impact of I110D versus I110E alone on the RuR sensitivity of K_2P_2.1 (TREK-1) (Figure 1E, Table 1) reinforces the idea that RuR molecular recognition requires both general electrostatic interactions and the direct coordination from the acidic sidechains. Taken together, our results indicate that details of these two factors tune the strength of the RuR interaction with the channel. Because of the largely conserved nature of the K_2P_ architecture in the region of the CAP and selectivity filter (Brohawn et al., 2012; Dong et al., 2015; Lolicato et al., 2017; Miller and Long, 2012), installing acidic residues simultaneously at the equivalents of the K_2P_2.1 (TREK-1) Ile110 and Asn147 positions should endow any K_2P_ of interest sensitive to nanomolar concentrations of RuR.

## Discussion

Polynuclear ruthenium compounds have been used for nearly 50 years to control the function of various ion channels (Arif Pavel et al., 2016; Braun et al., 2015; Caterina et al., 1999; Caterina et al., 1997; Chaudhuri et al., 2013; Coste et al., 2012; Czirjak and Enyedi, 2003; Dreses-Werringloer et al., 2013; Gonzalez et al., 2013; Guler et al., 2002; Kirichok et al., 2004; Ma, 1993; Ma et al., 2012; Moore, 1971; Musset et al., 2006; Rahamimoff and Alnaes, 1973; Smith et al., 1988; Story et al., 2003; Strotmann et al., 2000; Voets et al., 2004; Voets et al., 2002; Zhao et al., 2016). Yet, despite this widespread application in the study of a multitude of diverse ion channels, there is only limited visualization of how such compounds might interact with and affect the function of their targets (Choi et al., 2019). The K_2P_:RuR and K_2P_:Ru360 complexes presented here provide the first detailed structural views of how this class of inorganic polycations can inhibit ion channel function. The structures demonstrate general molecular recognition principles in which a channel uses acidic sidechains to coordinate both trinuclear, RuR, and dinuclear, Ru360, ruthenium amines through direct, multipronged electrostatic interactions. Both compounds block the flow of ions through the channel using a ‘finger in the dam’ mechanism that exploits the unique archway architecture that the K_2P_ CAP domain creates above the K_2P_ channel mouth (Figure 5A).

**Figure 5.**
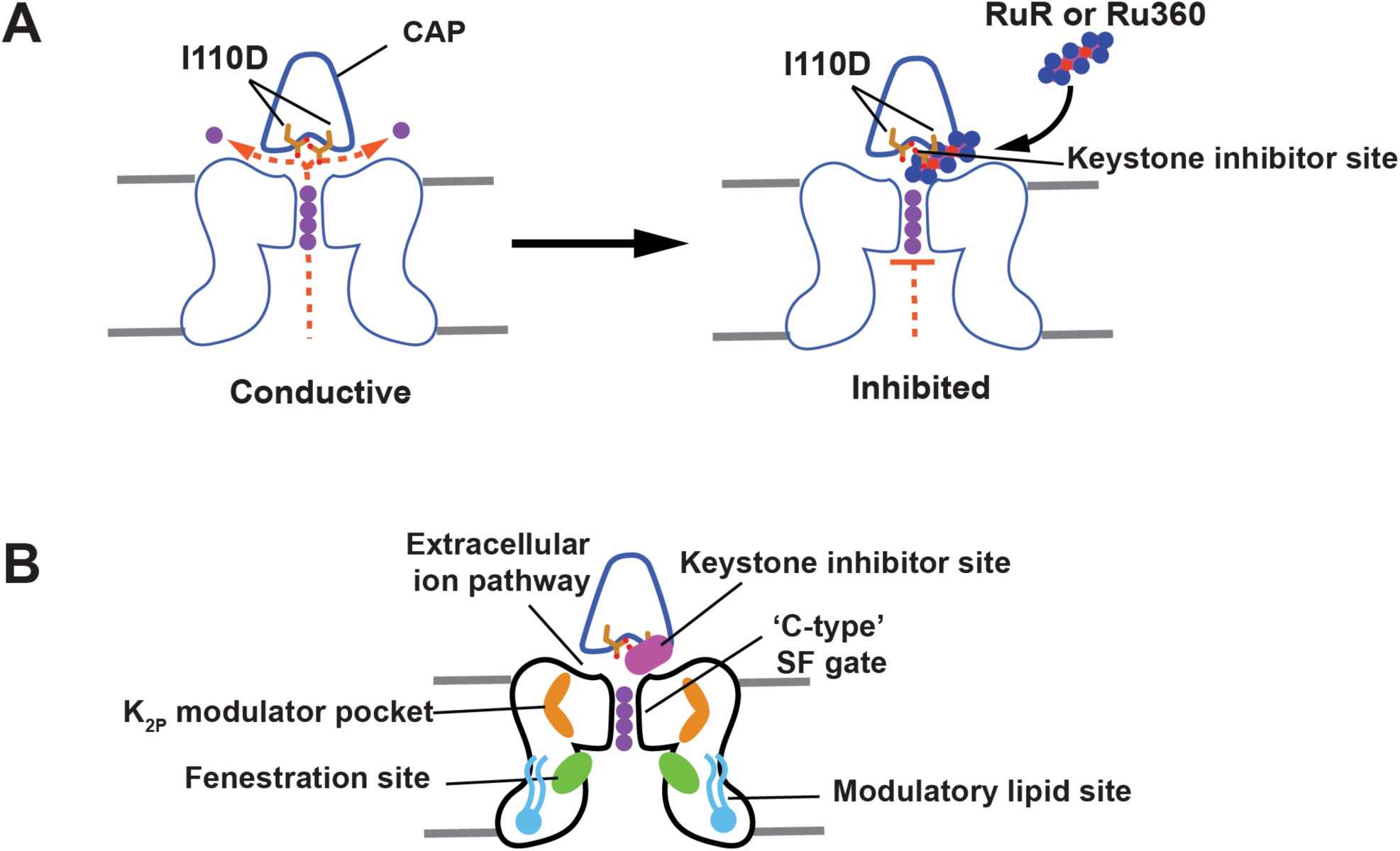
Mechanisms of small molecule K_2P_ modulation. (**A**) Cartoon diagram depicting the ‘finger in the dam’ mechanism of K_2P_ inhibition by polynuclear ruthenium amines. Ion flow is indicated by the dashed orange lines. Potassium ions are shown as purple circles. (**B**) Polysite model of K_2P_ modulation. Diagram shows structurally defined sites for K_2P_ modulators: Keystone inhibitor site (magenta), K_2P_ modulator pocket (orange) (Lolicato et al., 2017), fenestration site (green) (Dong et al., 2015; Schewe et al., 2019), and modulatory lipid site (cyan) (Lolicato et al., 2017). Extracellular ion pathway (EIP) and selectivity filter ‘C-type’ gate are indicated. CAP is outlined in blue and shows the position of the negatively charged residues required for the Keystone inhibitor site (sticks).

K_2P_s are the only potassium channel family that bears a CAP domain, an extracellular dimerization domain positioned directly above the channel pore (Brohawn et al., 2012; Miller and Long, 2012). The structures presented here show that a pair of aspartic acids located on the underside of the CAP archway at a site that controls the RuR response of natively-RuR sensitive K_2P_s, K_2P_10.1 (TREK-2) and K_2P_9.1 (TASK-3) (Czirjak and Enyedi, 2003; Gonzalez et al., 2013; Musset et al., 2006) and of the RuR-sensitive mutant K_2P_2.1 I110D (Figure 1E) (Braun et al., 2015), create a polycation binding site, the Keystone inhibitor site. This site forms the primary point of interaction with a single RuR or Ru360 that plugs one arm of the bifurcated EIP created by the CAP domain archway (Figs. 1E-F, 2E-F, 3G-H). This structural observation defines an unambiguous mechanism of action for how polyruthenium amines inhibit K_2P_s, as holding a large polycation above the channel pore would both physically block ion exit as well as provide an electrostatic barrier to permeant ion movement (Figure 5A). Ru360 a dinuclear oxo-bridged ruthenium amine inhibitor of the mitochondrial calcium uniporter (Baughman et al., 2011; Kirichok et al., 2004; Oxenoid et al., 2016) that had not previously been reported to affect potassium channels also inhibits engineered, K_2P_2.1 I110D, and natively-RuR sensitive K_2P_s, K_2P_10.1 (TREK-2), and K_2P_9.1 (TASK-3), in a manner similar to RuR (Figure 3E-F). Hence, even though RuR and Ru360 bind to a largely pre-formed binding site, there is sufficient plasticity to permit the binding of different types of polyruthenium cations. Consistent a binding site positioned above the selectivity filter and outside of the transmembrane electric field, polynuclear ruthenium amine inhibition of K_2P_s is independent of voltage (Figs. S1E-G, S4A-B, S5B-E and H-J) and the C-type gate activation (Figure 2D). Together, the data define a mechanism of action in which polynuclear ruthenium amines inhibit K_2P_ function by preventing ion flow out of the K_2P_ selectivity filter and through the EIP (Figure 5A).

Although many ion channels lack the extracellular archway made by the K_2P_ CAP domain, the structures of the K_2P_:polyruthenium amine complexes reveal general molecular recognition principles that are likely to be shared with RuR and Ru360 sensitive ion channels. The requirement to have a negatively charged residue at the Keystone inhibitor site for RuR (Figure 1E) (Braun et al., 2015; Czirjak and Enyedi, 2003; Gonzalez et al., 2013; Musset et al., 2006) and Ru360 (Figure 3F) responses, together with the observation that inhibitor potency is proportional to the total charge (Table 1) highlights the importance of electrostatic interactions for recognizing ruthenium polycations. The multipronged direct coordination of the RuR and Ru360 ruthenium amine moieties (Figure 1G, 2G, and 3G) shows the important role that direct coordination by acidic sidechains plays in binding both compounds. The observation that equivalently charged residues having different sidechain geometries, aspartate and glutamate, have differential effects on RuR potency at the Keystone inhibitor site (Figs. 1D and Table 1) highlights the importance of direct ligand coordination for tuning the strength of the interaction with polynuclear ruthenium amines. It should be noted that K_2P_4.1 (TRAAK) is inhibited by RuR but lacks a negatively charged residue in the Keystone inhibitor site (Figure S1A) and binds RuR with a higher stoichiometry than observed for the K_2P_ channels studied here (Table 1) (Braun et al., 2015; Czirjak and Enyedi, 2002). These differences suggest that RuR inhibition of K_2P_4.1 (TRAAK) occurs using a mechanism different from the ‘finger in the dam’ mechanism. Given the importance of direct interactions between RuR and Ru360 and the acidic sidechains in the target channels studied here and the fact that acidic residues are key to the RuR sensitivity of other channels (Zhao et al., 2018; Zhao et al., 2016), we expect that similar types of multipronged coordination by sidechain carboxylates are likely to contribute to the RuR and Ru360 block of other polynuclear ruthenium amine sensitive channels such as K_2P_4.1 (TRAAK), TRPs (Arif Pavel et al., 2016; Caterina et al., 1999; Caterina et al., 1997; Guler et al., 2002; Story et al., 2003; Strotmann et al., 2000; Voets et al., 2004; Voets et al., 2002), the mitochondrial calcium uniporter (MCU) (Chaudhuri et al., 2013; Kirichok et al., 2004; Moore, 1971; Rahamimoff and Alnaes, 1973), CALHM calcium channels (Choi et al., 2019; Dreses-Werringloer et al., 2013; Ma et al., 2012), ryanodine receptors (Ma, 1993; Smith et al., 1988), and Piezo channels (Coste et al., 2012; Zhao et al., 2016), even if the details of where the inhibitor binds to the channel differ.

K_2P_ channels can be modulated by a number of different small molecules and lipids (Sterbuleac, 2019). When placed in the context of previous structural studies of K_2P_ modulator interactions (Dong et al., 2015; Lolicato et al., 2017; Schewe et al., 2019), our studies highlight an emerging picture of the complex, multisite structural pharmacology that contributes to the control of K_2P_ function. The discovery of the Keystone inhibitor site reveals that there are at least four control sites that span from the inner leaflet of the bilayer to the extracellular parts of the channel through which exogenous molecules affect channel function (Figure 5B). These include a modulatory lipid binding site in the bilayer inner leaflet (Lolicato et al., 2017), the fenestration site residing at the intersection of the movable M4 transmembrane helix and the lower part of the selectivity filter that can be targeted by both small molecule activators and inhibitors (Dong et al., 2015; Schewe et al., 2019), the K_2P_ modulator pocket site (Lolicato et al., 2017), and the Keystone inhibitor site. Exploring the degree of conformational coupling among these modulatory sites will be important for understanding the extent of synergistic or antagonistic actions within the various classes of K_2P_ modulators. Acquiring this type of knowledge will be crucial for creating new interventions that could offer exquisite control of K_2P_ function.

The ‘finger in the dam’ inhibitory mechanism defined here provides a blueprint for the development of small molecule or protein-based K_2P_ modulators that could reach through the EIP to the Keystone inhibitor site. In this regard, designing compounds having moieties that interact with the Keystone inhibitor site but that also make contacts to non-conserved features of CAP exterior could yield subtype-selective modulators. Biologics, such as nanobodies, may be particularly suited to this type of molecular recognition mode.

Further, given the highly conserved nature of the K_2P_ channel architecture in the region of the CAP and selectivity filter (Brohawn et al., 2012; Dong et al., 2015; Lolicato et al., 2017; Miller and Long, 2012), the strategies we used to develop RuR super-responders should be applicable to other K_2P_ subfamily members to create subtypes endowed with RuR sensitivity. Such RuR-sensitive channels could be used to dissect the roles of various K_2P_s in their native physiological settings.

### Significance

Ruthenium Red (RuR) is a trinuclear, oxo-bridged ruthenium amine polycation that has many biological applications, including a ∼50 year legacy as an inhibitor of diverse classes of ion channels. RuR inhibits select members of the K_2P_ (KCNK) family, numerous TRP channels, the mitochondrial calcium uniporter, CALHM calcium channels, ryanodine receptors, and Piezo channels. Despite this remarkably wide range of ion cannel targets, there are extremely limited structural data describing how RuR binds to any ion channel target. Our studies show how two polyruthenium compounds, RuR and Ru360, inhibit K_2P_ channels through a ‘finger in the dam’ mechanism in which these polycations bind at a novel site, the ‘Keystone inhibitor site’, formed by acidic residue pair under the K_2P_ CAP domain archway above the channel pore. This series of structures, together with functional studies, outline the molecular recognition principles that govern how RuR and Ru360 bind to specific sites of proteins using a mixture of electrostatics and polyvalent coordination by acidic sidechains. These principles are likely to control RuR and Ru360 binding to a wide range of diverse ion channel targets. Moreover, we show that we can use knowledge of these factors to engineer RuR ‘super-responder’ K_2P_s that have RuR sensitivity in the low nanomolar range. The protein engineering strategy we define should be generally applicable to any K_2P_ of interest and provide a new method for dissecting the function of specific K_2P_s in complex settings such as neurons, the brain, and the cardiovascular system. Together, the data define a ‘finger in the dam’ inhibition mechanism acting at a novel K_2P_ inhibitor binding site. These findings highlight the polysite nature of K_2P_ pharmacology and provide a new framework for K_2P_ inhibitor development.

## Methods

### Molecular Biology

Murine K_2P_2.1 (TREK-1) (Gene ID 16526), K_2P_10.1 (TREK-2) (Gene ID: 72258), and K_2P_9.1 (TASK-3) (Gene ID: 223604) were each expressed from in a pGEMHE/pMO vector for two-electrode voltage clamp (TEVC) experiments as described previously (Lolicato et al., 2017; Pope et al., 2018). A previously described version of murine K_2P_2.1 (TREK-1) in a *Pichia pastoris* pPicZ plasmid, K_2P_2.1(TREK-1)_CRYST_ (Lolicato et al., 2017), encoding residues 21-322 and bearing the following mutations: K84R, Q85E, T86K, I88L, A89R, Q90A, A92P, N95S, S96D, T97Q, N119A, S300C, E306A, was used for structural studies. Mutants of K_2P_2.1 (TREK-1) and K_2P_2.1(TREK-1)_CRYST_ were generated using site-directed mutagenesis (PFU Turbo AD, Agilent) and verified by sequencing of the complete gene.

### Two-electrode voltage clamp (TEVC) electrophysiology

*Xenopus laevis* oocytes were harvested according to UCSF IACUC Protocol AN129690 and digested using collagenase (Worthington Biochemical Corporation, #LS004183, 0.7-0.8 mg mL^-1^) in Ca^2+^-free ND96 (96 mM NaCl, 2 mM KCl, 3.8 mM MgCl_2_, 5 mM HEPES pH 7.4) immediately post-harvest, as previously reported (Lolicato et al., 2017; Pope et al., 2018). Oocytes were maintained at 18°C in ND96 (96 mM NaCl, 2 mM KCl, 1.8 mM CaCl_2_, 2 mM MgCl_2_, 5 mM HEPES pH 7.4) supplemented with antibiotics (100 units mL^-1^ penicillin, 100 μg mL^-1^ streptomycin, 50 μg mL^-1^ gentimycin) and used for experiments within one week of harvest. mRNA for oocyte injection was prepared from plasmid DNA using mMessage Machine T7 Transcription Kit (Thermo Fisher Scientific), purified using RNEasy kit (Qiagen), and stored as stocks and dilutions in RNAse-free water at −80°C.

Defolliculated stage V-VI oocytes were microinjected with 50 nL of 0.1-6 ng mRNA and currents were recorded within 24-48 hours of injection. Oocytes were impaled by two standard microelectrodes (0.3-3.0 MΩ), filled with 3M KCl and subjected to constant perfusion of ND96. Currents were elicited from a −80 mV holding potential using a 500 ms ramp ranging from −140 to +50 mV.

Recording solutions containing Ruthenium red (RuR) (Millipore-Sigma, R2751) and Ru360 (Millipore-Sigma, Calbiochem – 557440) were prepared immediately prior to use. RuR was weighed and dissolved directly into ND96 at 200 μM in ND96 and then diluted into ND96 for tested experimental concentrations. The pH of the stock solution was checked to ensure no change occurred. Due to its instability in aqueous solutions, Ru360 solutions were covered with aluminum foil to minimize exposure to light and to avoid degradation. RuR and Ru360 were determined to be stable in recording solutions for duration of typical experiment length by measuring UV absorbance at 536 nm and 363 nm, respectively, before and after length of the recording session. ML335 was synthesized as described previously (Lolicato et al., 2017). ML335 recording solutions were prepared from a DMSO stock stored at −20°C (final DMSO concentration was 0.1%).

Data were recorded using a GeneClamp 500B (MDS Analytical Technologies) amplifier controlled by pClamp software (Molecular Devices), and digitized at 1 kHz using Digidata 1332A (MDS Analytical Technologies). For each recording, control solution (ND96) was perfused over a single oocyte until current was stable before switching to solutions containing the test compounds at various concentrations and again allowed to stabilize before recording final, stabilized trace. Fractional block at the potential of interest was determined as 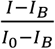 in which I is the measured current, I_0_ is the current in the absence of the test compound, and I_B_ is the basal current derived from an average of uninjected oocytes (n=14). For dose-response curves, each point is an average of at least three oocytes recorded from at least two independent batches of oocytes. Representative traces and dose response plots were generated in Graphpad Prism Version 5 (GraphPad Software, San Diego California USA, www.graphpad.com). Inhibition IC_50_s were estimated using an auto-fitted Hill equation with a Hill coefficient = 1.0.

### Protein expression

K_2P_2.1_CRYST_ I110D was expressed as a fusion protein having in series from the channel C-terminus a 3C protease site, green fluorescent protein (GFP), and His_10_ tag as described previously for K_2P_2.1_cryst_ (Lolicato et al., 2017). Linearized plasmid DNA (*PmeI*) was introduced into *Pichia pastoris* strain SMD1163H via electroporation. Strains with highest incorporation were selected for on YPD plates containing 1-2 mg mL^-1^ zeocin. Individual colonies were screened using fluorescence size exclusion chromatography (FSEC) (Kawate and Gouaux, 2006) to identify strain with highest expression level as described previously (Lolicato et al., 2017). The best FSEC candidate was used to inoculate a starter culture (60-120 mL) in minimal media (1% glycerol, 100 mM potassium phosphate pH 6.0, 0.4 mg L^-1^ biotin, 1X YNB from Invitrogen) supplemented with 1 mg mL^-1^ zeocin and cultured in shaker flask for 2 days at 29^°^C. The starter culture was then used to inoculate a large scale (6-12L) culture in shaker flasks containing minimal media without zeocin. Cells were grown at 29^°^C over two days in minimal media containing 1% glycerol. Cells were centrifuged at 3000g (6 min, 20^°^C) and pellet was resuspended in minimal induction media (100 mM potassium phosphate pH 6.0, 0.4 mg L^-1^ biotin, 1X YNB) containing 0.5% methanol. After 24 hrs, 0.5% methanol was added to each flask. Cells were harvested (6000g, 20 min, 4^°^C) two days after induction, snap-frozen in liquid nitrogen, and stored at −80°C.

### Protein Purification

Purified K_2P_2.1_cryst_ I110D was obtained from preparations using 100-200g of cell mass cryo-milled (Retsch, MM400) in liquid nitrogen (5 × 3 min, 25 Hz). All purification steps were carried out at 4°C and purification conditions were similar to those previously reported for K_2P_2.1_cryst_ (Lolicato et al., 2017). Cell powder was solubilized at a ratio of 3 grams of cells per mL of lysis buffer containing 200 mM KCl, 21 mM octyl glucose neopentyl glycol (OGNG, Anatrace), 30 mM *n*-heptyl-β-D-thioglucopyranoside (HTG, Anatrace), 0.1% cholesterol hemisuccinate (CHS, Anatrace), 100 mM Tris-Cl, pH 8.2, 1 mM PMSF and 0.1 mg/mL DNaseI. Following 3 hour membrane solubilization, the sample was centrifuged at 40,000g for 45 min at 4°C. After centrifugation, supernatant was incubated with anti-GFP nanobodies immobilized on CNBr-activated sepharose resin (GE Healthcare, 17-0430-01) at a ratio of 1 mL resin per 10 g of cells and gently rotated on an orbital rocker for 3 hours. Resin was collected in a gravity column (Econo-Pac, 1.5 × 12 cm, BioRad) and washed with 10 CV each, buffers A-C (A-C: 200 mM KCl, 50 mM Tris-Cl pH 8.0, 15 mM HTG; A: 10 mM OGNG, 0.018% CHS; B: 5 mM OGNG, 0.018% CHS; C: 3.85 mM OGNG, 0.0156% CHS), applied in series to reduce the detergent concentration and wash away cell debris (30 CV total). The GFP-affinity tag was cleaved overnight on column using two CV of buffer C supplemented with 350 mM KCl, 1 mM EDTA and 3C protease (Shaya et al., 2011). Cleaved protein was eluted from resin with two CV of size-exclusion buffer (SEC: 200 mM KCl, 20 mM Tris-Cl pH 8.0, 2.1 mM OGNG, 15 mM HTG, 0.012% CHS), concentrated and applied to a Superdex 200 (GE, 10/300) pre-equilibrated with SEC buffer. Peak fractions were evaluated by SDS-PAGE (15% acrylamide) for purity, pooled and concentrated for crystallization.

### Crystallization, structure determination, and refinement

Purified K_2P_2.1_cryst_-I110D was concentrated to 6 mg mL^-1^ before crystallization using hanging-drop vapor diffusion at 4°C using a mixture of 0.2 µL protein to 0.1 µL reservoir solution, over 100 µL reservoir of 20-25% PEG400, 200 mM KCl, 100 mM HEPES pH 8.0 or 7.1 and 1-2 mM CdCl_2_. Crystals appeared within 1-2 days and grew to full size within 2 weeks. Crystals were cryoprotected in solution containing 200 mM KCl, 0.2% OGNG, 15 mM HTG, 0.02% CHS, 100 mM HEPES pH 8.0 or 7.1 and 1-2 mM CdCl_2_, with 5% increase in PEG400 up to final concentration of 38% before flash freezing in liquid nitrogen. For compound bound structures, cryoprotected crystals were also soaked in final cryoprotection solution containing 1 mM each RuRed, RuRed+ML335 or Ru360, sourced as described in TEVC methods, for at least 1 hour prior to flash freezing in liquid nitrogen.

Datasets were collected at 100 K using synchrotron radiation at APS GM/CAT beamline 23-IDB/D Chicago, Illinois using a wavelength of 1.0332 Å, processed with XDS (Kabsch, 2010) and scaled and merged with Aimless (Evans and Murshudov, 2013). Highest resolution structures were obtained from crystals that were soaked with the ruthenium compounds. Structure determination of low resolution datasets from complexes obtained by co-crystallization indicated that there was no difference in the RuR position in complexes made by either soaking or co-crystallization. Final resolution cutoff was 3.40 Å, 3.49 Å, 3.00 Å and 3.51 Å for K_2P_2.1_cryst_ I110D, K_2P_2.1_cryst_ I110D:RuR, K_2P_2.1_cryst_ I110D:RuR:ML335 and K_2P_2.1_cryst_ I110D:Ru360 structures, respectively, using the CC_1/2_ criterion (Diederichs and Karplus, 2013). K_2P_2.1_cryst_-I110D was solved by molecular replacement utilizing K_2P_4.1 (G124I) structure (PDB: 4RUE) as search model. For compound bound structures, the K_2P_2.1_cryst_-I110D model was used as the molecular replacement search model. Electron density maps were improved through several cycles of manual model rebuilding, using COOT (Emsley and Cowtan, 2004), REFMAC (CCP4), and PHENIX (Adams et al., 2010).

### Mutant cycle analysis

Double mutant cycle analysis (Carter et al., 1984; Hidalgo and MacKinnon, 1995) was carried out using the equation 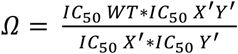 in which Ω is the coupling factor (Hidalgo and MacKinnon, 1995) and IC_50_WT, IC_50_X’, IC_50_Y’ and IC_50_X’Y’ are the IC_50_ values for the wild-type, each single mutant of the X-Y pair, and the double mutant, respectively. As wild type K_2P_2.1 (TREK-1) is unaffected by RuR, the free energy of the interaction is zero and hence, K_a_ and K_d_ = 1. Coupling energy, ΔΔG_Ω_, was calculated as ΔΔ*G*_*Ω*_ = *RTLnΩ* where R=1.987 cal mol^-1^ deg^-1^ and T= 298K.

## Acknowledgements

We thank V. Nguyen for help in the initial stages of this project, Z. Wong for technical assistance, A. Natale for help with structure comparisons, M. Grabe and for helpful discussions, and Y. Kirichok and members of the Minor lab for comments on the manuscript. This work was supported by grant NIH-R01-MH093603 to D.L.M.

## Author Contributions

L.P. and D.L.M. conceived the study and designed the experiments. L.P. performed molecular biology experiments, two-electrode voltage clamp recordings, expressed, purified, and crystallized the proteins, and collected diffraction data. L.P. and M.L. determined the structures. L.P. and M.L. analyzed the data. D.L.M. analyzed data and provided guidance and support. L.P., M.L., and D.L.M. wrote the paper.

## Data and materials availability

Coordinates and structures factors for the structures of K_2P_2.1 I110D (6V36), K_2P_2.1 I110D:RuR (6V3I), K_2P_2.1 I110D:RuR:ML335 (6V37), and K_2P_2.1 I110D:Ru360 (6V3C) are deposited with RCSB and will be released immediately upon publication.

