## Supplementary Figures and Tables for "Polynuclear ruthenium amines inhibit K_2P_ channels via a ‘finger in the dam’ mechanism"

2-Dec-19

**Short Title:** $K_{2P}$ inhibition by polynuclear ruthenium amines

Lianne Pope[1], Marco Lolicato[1#], and Daniel L. Minor, Jr.[1-5*]

[1]Cardiovascular Research Institute
[2]Departments of Biochemistry and Biophysics, and Cellular and Molecular Pharmacology
[3]California Institute for Quantitative Biomedical Research
[4]Kavli Institute for Fundamental Neuroscience
University of California, San Francisco, California 93858-2330 USA

[5]Molecular Biophysics and Integrated Bio-imaging Division
Lawrence Berkeley National Laboratory, Berkeley, CA 94720 USA

#Current address:
Department of Molecular Medicine
University of Pavia
Pavia ITALY

**Keywords:** $K_{2P}$ channel, X-ray crystallography, Ruthenium Red, Ru360, electrophysiology, Keystone inhibitor site

**Supplementary Materials**

**Table S1** Crystallographic data collection and refinement statistics.

**Table S2** Structure comparison of RuR and Ru360 complexes.

**Figure S1** $K_{2P}$ CAP sequences and $K_{2P}$2.1 CAP mutant functional properties.

**Figure S2** $K_{2P}$2.1 I110D structures and structure comparisons.

**Figure S3** $K_{2P}$2.1 I110D ML335 response and structure of the $K_{2P}$2.1 I110D:RuR:ML335 complex.

**Figure S4** Ru360 inhibits $K_{2P}$ channels.

**Figure S5** $K_{2P}$ selectivity filter (SF) sequences and RuR responses of $K_{2P}$2.1 SF mutants.

**Figure S6** Double mutant cycle analysis.

**Movie S1** RuR block of $K_{2P}$ channels.

### Figure S1

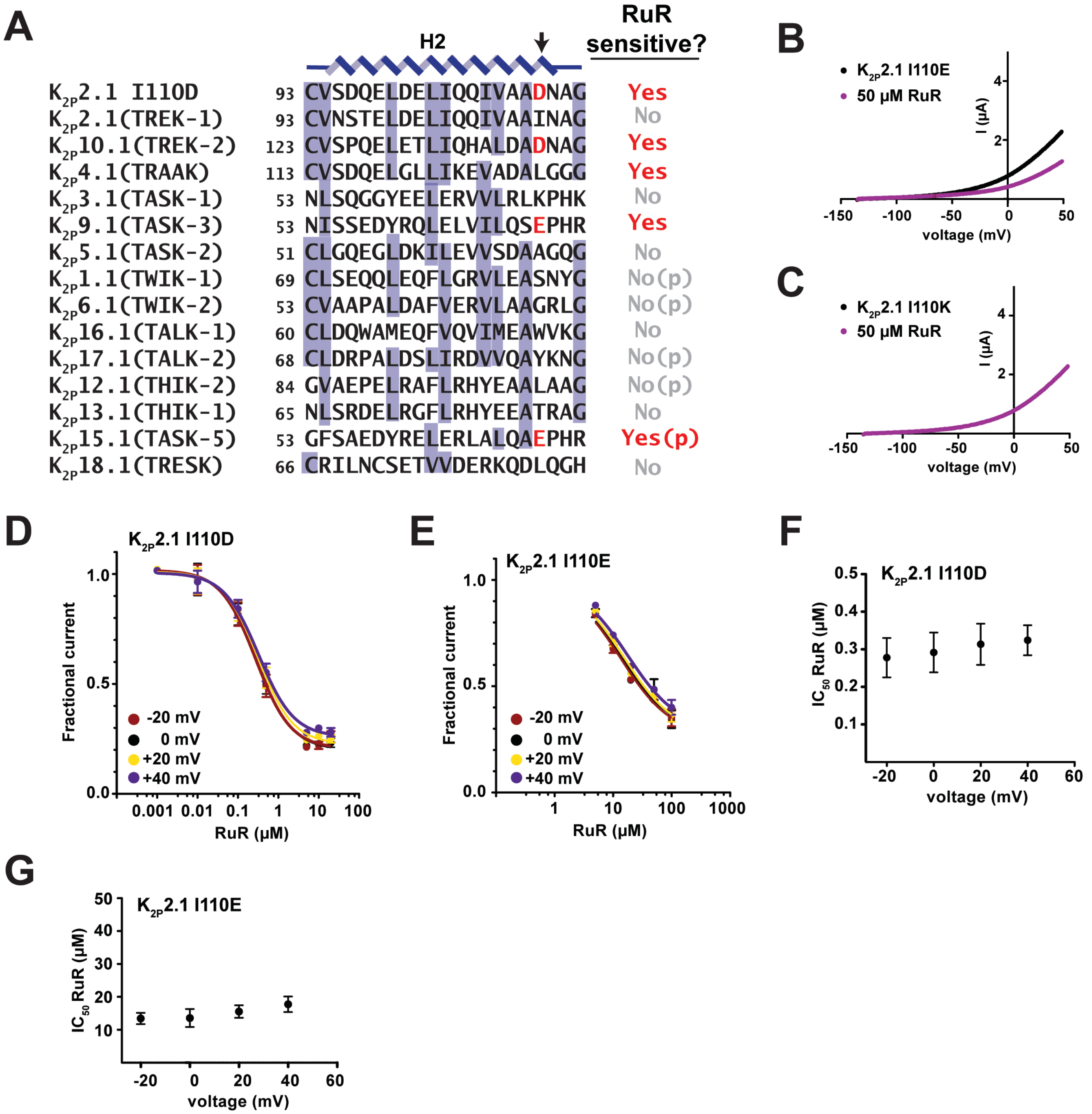

**Figure S1 $K_{2P}$ CAP sequences and $K_{2P}$2.1 CAP mutant functional properties.** (**A**) CAP H2 helix sequences for the indicated $K_{2P}$ channels. Measured (Braun et al., 2015; Czirjak and Enyedi, 2003; Gonzalez et al., 2013; Musset et al., 2006) or predicted (p) Ruthenium Red (RuR) sensitivity is indicated. Arrow indicates the position of the RuR sensitivity determinant (amino acids highlighted in red). $K_{2P}$2.1 I110D , $K_{2P}$2.1 (TREK-1) AAD47569.1, $K_{2P}$10.1 (TREK-2) BAF83207, $K_{2P}$4.1 (TRAAK) AAI10328.1, $K_{2P}$3.1 (TASK-1) NP_002237.1, $K_{2P}$9.1 (TASK-3) NP_001269463.1, $K_{2P}$5.1 (TASK-2) NP_003731.1, $K_{2P}$1.1 (TWIK-1) NP_002236.1, $K_{2P}$6.1 (TWIK-2) NP_004814.1, $K_{2P}$16.1 (TALK-1) NP_001128577.1, $K_{2P}$17.1 (TALK-2) AAK28551.1, $K_{2P}$12.1 (THIK-2) NP_071338.1, $K_{2P}$13.1 (THIK-1) NP_071337.2,

$K_{2P}$15.1 (TASK-5) EAW75900.1, $K_{2P}$18.1 (TRESK) NP_862823.1. $K_{2P}$15.1 (TASK-5) cannot be functionally expressed (Enyedi and Czirjak, 2010). (**B**) and (**C**) Exemplar TEVC recordings of (**B**) $K_{2P}$2.1 I110E and (**C**) $K_{2P}$2.1 I110K responses to 50 μM RuR (magenta). (**D-G**) Analysis of the voltage-dependence of RuR inhibition of $K_{2P}$2.1 I110D and $K_{2P}$2.1 I110E. (**D**) and (**E**) Dose-response curves at -20 mV (red), 0 mV (black), +20 mV (yellow), and +40 mv (purple) for (**D**) $K_{2P}$2.1 I110D and (**E**) $K_{2P}$2.1 I110E. (**F**) and (**G**) RuR $IC_{50}$ (μM) as a function of voltage for (**D**) $K_{2P}$2.1 I110D and (**E**) $K_{2P}$2.1 I110E. Error bars are SEM.

### Figure S2

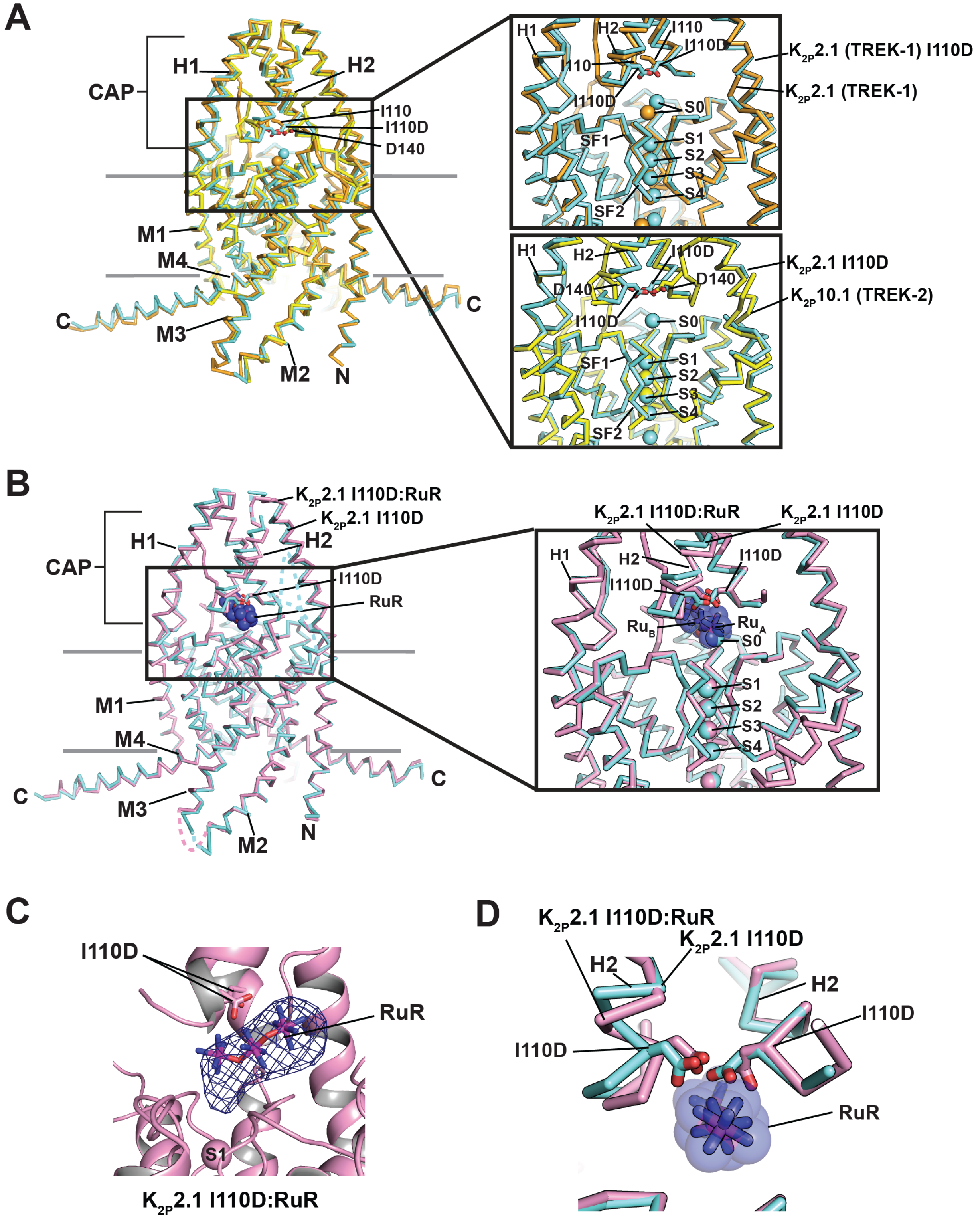

**Figure S2 $K_{2P}$2.1 I110D structures and structure comparisons.** (**A**) Superposition of $K_{2P}$2.1 (TREK-1) (orange) (PDB:6CQ6)(Lolicato et al., 2017), $K_{2P}$2.1 I110D (cyan), and $K_{2P}$10.1 (TREK-2) (PDB:4BW5) (yellow)(Dong et al., 2015). Insets show comparison of area around the I110D mutation. Top, $K_{2P}$2.1 (TREK-1) and $K_{2P}$2.1 I110D; Bottom $K_{2P}$2.1 I110D and $K_{2P}$10.1 (TREK-2). (**B**) Superposition of

$K_{2P}$2.1 I110D (cyan) and $K_{2P}$2.1 I110D:RuR (pink). Inset shows area around the Keystone inhibitor site. RuR is shown as sticks with a semi-transparent surface. (**C**) Exemplar Fo-Fc density (3$\sigma$)(dark blue) for the $K_{2P}$2.1 I110D:RuR complex (pink). RuR is shown as sticks. (**D**) Close up view of $K_{2P}$2.1 I110D (cyan) and $K_{2P}$2.1 I110D:RuR (pink) showing conformational changes in the Keystone inhibitor site. RuR is shown as sticks with a semi-transparent surface.

### Figure S3

Pope *et al.*

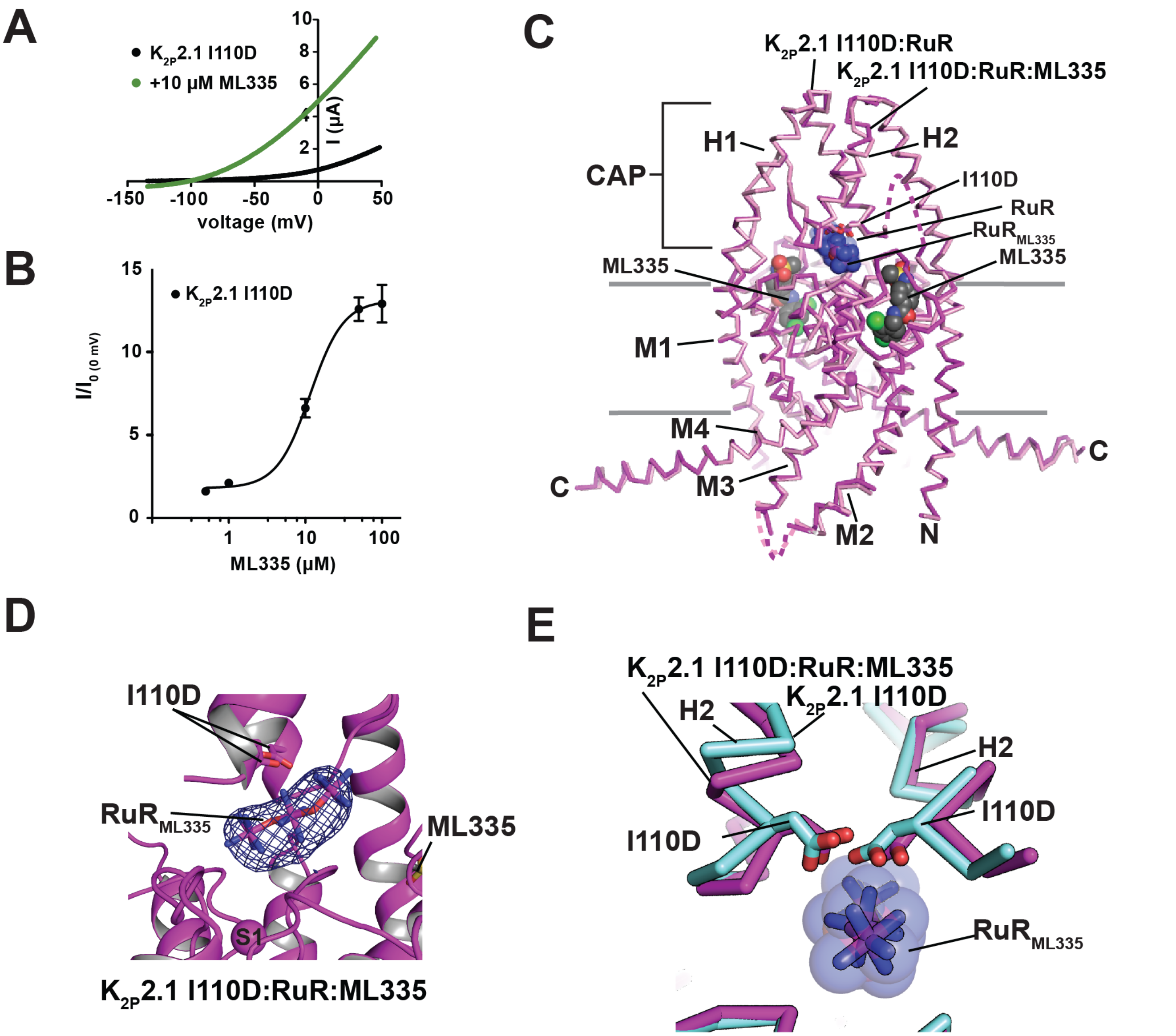

**Figure S3 $K_{2P}$2.1 I110D ML335 response and structure of the $K_{2P}$2.1 I110D:RuR:ML335 complex.** (**A**) Exemplar two-electrode voltage clamp recordings of the response of $K_{2P}$2.1 I110D (black) to 10 μM ML335 (green). (**B**) ML335 dose-response curve for $K_{2P}$2.1 I110D. $EC_{50}$ = 11.8 ± 2.3 μM, matching that of $K_{2P}$2.1 (TREK-1) (14.3 ± 2.7μM (Lolicato et al., 2017)). (**C**) Superposition of $K_{2P}$2.1 I110D:RuR (pink) and $K_{2P}$2.1 I110D:RuR:ML335 (magenta). I110D is shown as sticks. RuR and ML335 are shown in space filling representation. (**D**) Exemplar Fo-Fc density (3σ)(dark blue) for the $K_{2P}$2.1 I110D:RuR:ML335 complex (magenta), and (I110D, RuR, Ru360, and ML335 are shown as sticks. S1 selectivity filter ion is labeled. (**E**) Close up view of $K_{2P}$2.1 I110D (cyan) and $K_{2P}$2.1 I110D:RuR:ML335 (magenta) CAP base.

### Figure S4

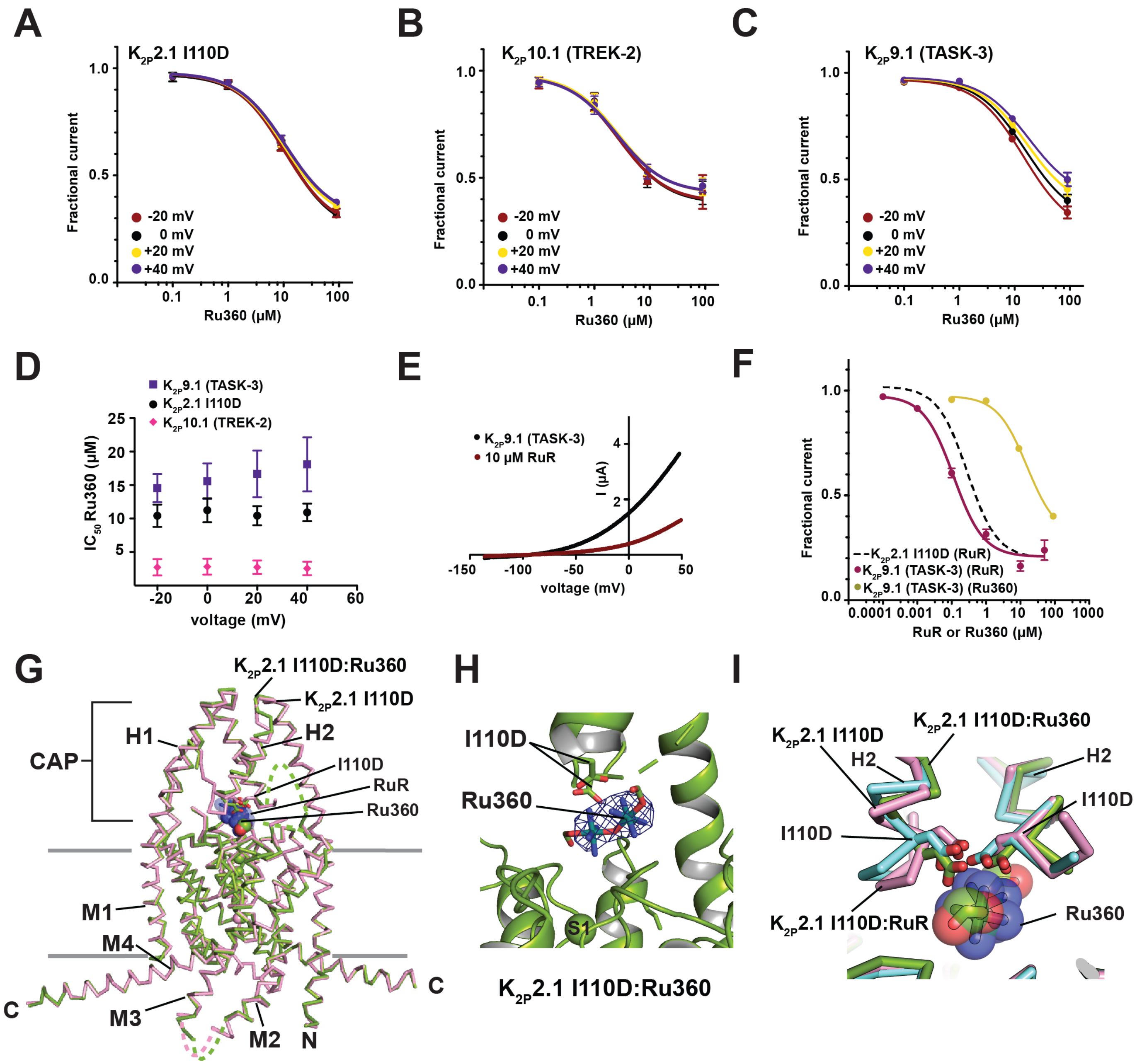

**Figure S4 Ru360 inhibits K2P channels.** (**A-C**) Ru360 dose-response curves at -20 mV (red), 0 mV (black), +20 mV (yellow), and +40 mv (purple) for (**A**) K2P2.1 I110D, (**B**) K2P10.1 (TREK-2), and (**C**) K2P9.1 (TASK-3). (**D**) RuR $IC_{50}$ (μM) voltage-dependence for K2P2.1 I110D (black circles), K2P10.1 (TREK-2) (pink diamonds), and K2P9.1 (TASK-3) (purple squares). (**E**) Exemplar TEVC recordings of K2P9.1 (TASK-3) alone (black) and in the presence of 10 μM RuR (dark red). (**F**) K2P9.1 (TASK-3) dose-response curves for RuR (dark red) and Ru360 (olive) (from Figure 3F). Dashed line shows RuR dose-response for K2P2.1 I110D from Figure 1D. (**G**) Superposition of K2P2.1 I110D:RuR (pink) and K2P2.1 I110D:Ru360 (green). I110D is shown as sticks. (**H**) Exemplar Fo-Fc density (3σ)(dark blue) for the K2P2.1 I110D:Ru360 complex (green). I110D and Ru360 are shown as sticks. S1 selectivity filter ion is labeled. (**I**) Close up view of $K_{2P}$2.1 I110D:Ru360 (green). $K_{2P}$2.1 I110D (cyan), and $K_{2P}$2.1 I110D:RuR (pink) CAP base. RuR and Ru360 and are shown in space filling representation.

### Figure S5

Pope *et al.*

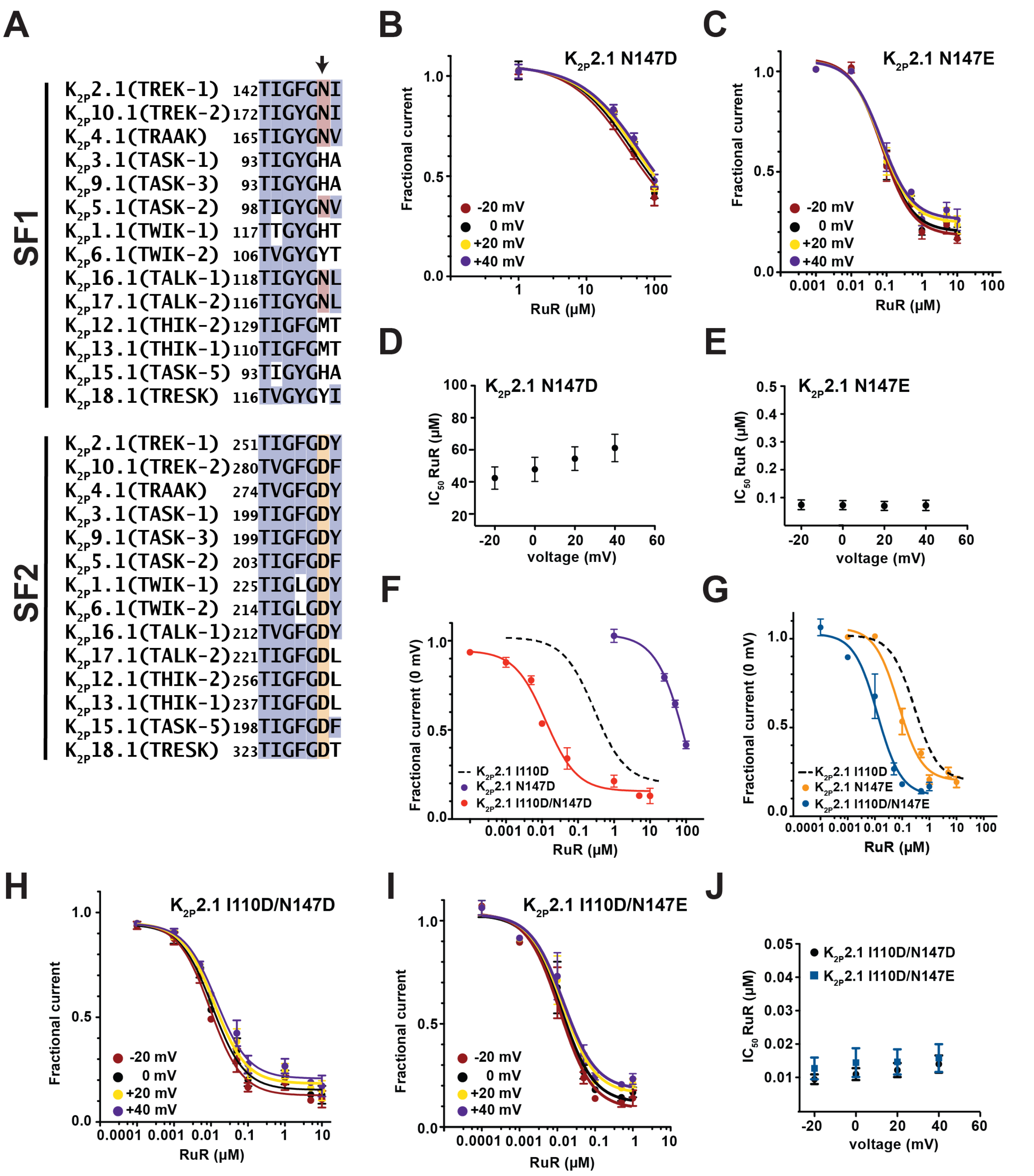

**Figure S5 $K_{2P}$ selectivity filter (SF) sequences and RuR responses of $K_{2P}$2.1 SF mutants.** (**A**) Sequence alignment of the selectivity filter 1 (SF1) and selectivity filter 2 (SF2) sequences of the following human K2P channels. $K_{2P}$2.1 (TREK-1) AAD47569.1, $K_{2P}$10.1 (TREK-2) BAF83207, $K_{2P}$4.1 (TRAAK) AAI10328.1, $K_{2P}$3.1 (TASK-1) NP_002237.1, $K_{2P}$9.1 (TASK-3) NP_001269463.1, $K_{2P}$5.1 (TASK-2) NP_003731.1, $K_{2P}$1.1 (TWIK-1) NP_002236.1, $K_{2P}$6.1 (TWIK-2) NP_004814.1, $K_{2P}$16.1 (TALK-1)

NP_001128577.1, $K_{2P}$17.1 (TALK-2) AAK28551.1, $K_{2P}$12.1 (THIK-2) NP_071338.1, $K_{2P}$13.1 (THIK-1) NP_071337.2, $K_{2P}$15.1 (TASK-5) EAW75900.1, $K_{2P}$18.1 (TRESK) NP_862823.1. SF1 and SF2 sequence and numbers for $K_{2P}$2.1 (TREK-1)$_{cryst}$ (PDB:6CQ6)(Lolicato et al., 2017) are identical to that of $K_{2P}$2.1 (TREK-1) AAD47569.1. (**B-C**) RuR dose-response curves at -20 mV (red), 0 mV (black), +20 mV (yellow), and +40 mv (purple) for (**B**) $K_{2P}$2.1 N147D and (**C**) $K_{2P}$2.1 N147E. (**D-E**) $IC_{50}$ voltage dependence for (**D**) $K_{2P}$2.1 N147D and (**E**) $K_{2P}$2.1 N147E. (**F**-**G**) RuR dose-response curves for (**F**) $K_{2P}$2.1 N147D (purple) and $K_{2P}$2.1 I110D/N147D (red) and (**G**) $K_{2P}$2.1 I110E (orange) and $K_{2P}$2.1 I110D/N147E (blue). Dashed lines show RuR response of $K_{2P}$2.1 I110D from Figure 1D. (**H-I**) RuR dose-response curves at -20 mV (red), 0 mV (black), +20 mV (yellow), and +40 mv (purple) for (**H**) $K_{2P}$2.1 I110D/N147D and (**I**) $K_{2P}$2.1 I110D/N147E. (**J**) $IC_{50}$ voltage dependence for $K_{2P}$2.1 I110D/N147D (black circles) and $K_{2P}$2.1 I110D/N147E (blue squares). Error bars are SEM.

### Figure S6

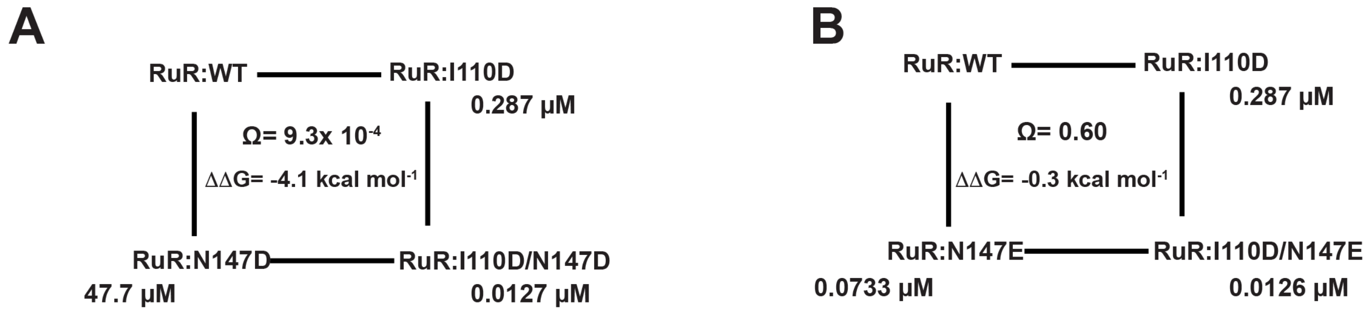

**Figure S6 Double mutant cycle analysis.** Double mutant cycle analysis (Carter et al., 1984; Hidalgo and MacKinnon, 1995) for the RuR responses of (**A**) $K_{2P}$2.1 I110D/N147D and (**B**) $K_{2P}$2.1 I110D/N147E. $\Omega = \frac{IC_{50}\ X'Y'}{IC_{50}\ X' * IC_{50}\ Y'}$. as $\Delta\Delta G_{\Omega} = RTLn\Omega$ where R=1.987 cal mol$^{-1}$ deg$^{-1}$ and T= 298K.

**Movie S1 RuR block of $K_{2P}$ channels.** Morph between the $K_{2P}$2.1 I110D and $K_{2P}$2.1 I110D:RuR complex structures. I110D sidechains are shown as sticks. RuR is shown in space filling.

**Table S1 Data collection and refinement statistics**

| | $K_{2P}$2.1 (TREK-1) I110D (PDB:6V36) | $K_{2P}$2.1 (TREK-1) I110D:RuR (PDB:6V3I) | $K_{2P}$2.1 (TREK-1) I110D:RuR:ML335 (PDB: 6V37) | $K_{2P}$2.1 (TREK-1) I110D:Ru360 (PDB:6V3C) |
|---|---|---|---|---|
| **Data collection** | | | | |
| Space group | $P2_12_12_1$ | $P2_12_12_1$ | $P2_12_12_1$ | $P2_12_12_1$ |
| Cell dimensions | | | | |
| *a*, *b*, *c* (Å) | 69.19/120.40/128.35 | 67.95/120.3/127.82 | 67.02/118.74/129.04 | 67.74/120.71/127.27 |
| $\alpha, \beta, \gamma$ (°) | 90/90/90 | 90/90/90 | 90/90/90 | 90/90/90 |
| Resolution (Å) | 46.7 – 3.40 (3.67 - 3.40) | 46.6 - 3.49 (3.82 - 3.49) | 46.5 - 3.00 (3.18 - 3.00) | 46.4 - 3.51 (3.85 - 3.51) |
| $R_{merge}$ | 0.072 (5.87) | 0.089 (3.80) | 0.085 (2.45) | 0.110 (8.64) |
| $I/\sigma(I)$ | 9.6 (0.5) | 9.8 (0.8) | 16.8 (1.2) | 10.8 (0.4) |
| $CC_{1/2}$ | 0.999 (0.128) | 1.000 (0.331) | 0.996 (0.553) | 1.000 (0.149) |
| Completeness (%) | 99.9 (100.0) | 100.0 (100.0) | 100.0 (100.0) | 100.0 (100.0) |
| Redundancy | 6.5 (6.7) | 8.8 (9.0) | 13.2 (13.7) | 13.3 (13.7) |
| **Refinement** | | | | |
| Resolution (Å) | 15.00 - 3.40 | 15.00 – 3.49 | 14.99 – 3.00 | 14.98 – 3.51 |
| No. reflections | 14468 | 13484 | 21051 | 12595 |
| $R_{work}$ / $R_{free}$ | 28.6/31.6 | 26.5/32.7 | 26.7/31.4 | 31.0/32.6 |
| No. atoms | 4124 | 4192 | 4309 | 4063 |
| Protein | 4066 | 4092 | 4180 | 3993 |
| Ligand/ion | 55 | 98 | 127 | 68 |
| $K^+$ | 6 | 5 | 5 | 5 |
| $Cd^{++}$ | 3 | 3 | 3 | 3 |
| Lipid | 46 | 63 | 82 | 43 |
| ML335 | 0 | 0 | 1 | 0 |
| RuR | 0 | 1 | 1 | 0 |
| Ru360 | 0 | 0 | 0 | 1 |
| Water | 3 | 2 | 2 | 2 |
| *B* factors | | | | |
| Protein | 227.94 | 191.96 | 145.60 | 200.63 |
| Ligand/ion | 195.40 | 196.35 | 150.32 | 182.87 |
| R.M.S. deviations | | | | |
| Bond lengths (Å) | 0.004 | 0.004 | 0.003 | 0.002 |
| Bond angles (°) | 0.913 | 0.760 | 0.628 | 0.559 |
| Ramachandran | | | | |
| Favored (%) | 94.6 | 93.9 | 92.5 | 96.6 |
| Allowed (%) | 4.8 | 5.1 | 6.7 | 3.4 |
| Outliers (%) | 0.6 | 1.0 | 0.8 | 0.0 |

Each dataset was derived from a single crystal.

[a] Values in parentheses are for highest-resolution shell.

**Table S2 Structure Comparisons of RuR and Ru360 complexes**

| | | | RMSD (Å) |
|---|---|---|---|
| $K_{2P}$2.1 I110D | | $K_{2P}$2.1 (TREK-1) (6CQ6)(Lolicato et al., 2017) | 0.575 |
| $K_{2P}$2.1 I110D | | $K_{2P}$10.1 (TREK-2) (4BW5)(Dong et al., 2015) | 0.938 |
| $K_{2P}$2.1 I110D | versus | $K_{2P}$2.1 I110D:RuR | 0.688 |
| $K_{2P}$2.1 I110D | | $K_{2P}$2.1 I110D:Ru360 | 0.665 |
| $K_{2P}$2.1 I110D:RuR:ML335 | | $K_{2P}$2.1 I110D:RuR | 0.507 |
| $K_{2P}$2.1 I110D:RuR:ML335 | | $K_{2P}$2.1 (TREK-1):ML335 (6CQ8)(Lolicato et al., 2017) | 0.480 |
| $K_{2P}$2.1 I110D:Ru360 | | $K_{2P}$2.1 I110D:RuR | 0.561 |

RMSDs were calculated using C$\alpha$ atoms in the specified residue ranges after optimal translational and rotational alignment of those atoms. Calculations were performed with the MDAnalysis python package version 0.19.2 (Michaud-Agrawal et al., 2011). Residue ranges include all of the $\alpha$-helical portions of the structures as follows: $K_{2P}$2.1 (TREK-1): 45-93, 96-110, 126-190, 206-259, 268-300; $K_{2P}$10.1 (TREK-2): 75-123, 126-140, 156-220, 236-289, 299-331. 4BW5 comparison is for chains A and B from the asymmetric unit.
